# The relationship between event boundary strength and pattern shifts across the cortical hierarchy during naturalistic movie-viewing

**DOI:** 10.1101/2024.04.10.588931

**Authors:** Yoonjung Lee, Janice Chen

**Affiliations:** Department of Psychological and Brain Sciences, Johns Hopkins University, Baltimore, MD 21218, USA

**Keywords:** event boundary, cortical hierarchy, timescales, default mode network, hippocampus

## Abstract

Our continuous experience is spontaneously segmented by the brain into discrete events. However, the beginning of a new event (an event boundary) is not always sharply identifiable: phenomenologically, event boundaries vary in salience. How are the response profiles of cortical areas at event boundaries modulated by boundary strength during complex, naturalistic movie-viewing? Do cortical responses scale in a graded manner with boundary strength, or do they merely detect boundaries in a binary fashion? We measured “cortical boundary shifts” as transient changes in multi-voxel patterns at event boundaries with different strengths (weak, moderate, and strong), determined by across-subject agreement. Cortical regions with different processing timescales were examined. In auditory areas, which have short timescales, cortical boundary shifts exhibited a clearly graded profile both in group-level and individual-level analyses. In cortical areas with long timescales, including the default mode network, boundary strength modulated pattern shift magnitude at the individual subject level. We also observed a positive relationship between boundary strength and the extent of temporal alignment of boundary shifts across different levels of the cortical hierarchy. Additionally, hippocampal activity was highest at event boundaries for which cortical boundary shifts were most aligned across hierarchical levels. Overall, we found that event boundary strength modulated cortical pattern shifts strongly in sensory areas and more weakly in higher-level areas, and that stronger boundaries were associated with greater alignment of these shifts across the cortical hierarchy.

## Introduction

In the human brain, our continuous experience is segmented into meaningful units or events; moments at which one segment is perceived to end and a new one begins are referred to as event boundaries. Event segmentation theory (EST) proposes that an active mental model about “what is happening now” (i.e., an event model) is updated at event boundaries—when the brain fails to predict what will happen in the immediate future (Zacks et al., 2007). Considerable evidence suggests that perceiving event boundaries influences memory organization. Items encoded within a stable context are more strongly bound together in long-term memory representations than those which occur in different contexts, irrespective of their absolute distance in time (DuBrow & Davachi, 2013, 2016; Ezzyat & Davachi, 2011, 2014; Heusser et al., 2018); event boundaries seem to act as access points when retrieving memories of naturalistic experiences (Michelmann et al., 2023). Furthermore, during naturalistic movie-viewing, hippocampal activity at the offset of events is associated with later retrieval success or the strength of event memory reactivation (Baldassano et al., 2017; Ben-Yakov & Dudai, 2011; Ben-Yakov et al., 2013).

Phenomenologically, people experience a spectrum of event boundary strengths. In narrative comprehension studies, sudden changes in spatial or temporal context (e.g., a temporal shift to “an hour later”) are commonly regarded as strong transitions (for review, Zwaan et al., 1995). Conversely, small changes, such as a minimal temporal shift to “a moment later”, are generally perceived to be weak or barely detectable transitions (Zwaan, 1996). Additionally, event segmentation can be performed at different levels of granularity (coarse-grained vs. fine-grained), and such procedures reveal a hierarchical structure: coarse-grained events are divided into smaller segments by fine-grained event boundaries, and some event boundaries in the fine-grained set temporally align with those in the coarse-grained set (Zacks, Tversky, et al., 2001). Thus, observations from prior studies indicate the existence of salient, easily identifiable event boundaries that individuals reliably recognize when asked to perform event segmentation at any level of coarseness.

Interestingly, recent research suggests that hippocampal responses at movie event boundaries show a graded profile, wherein greater univariate activity is associated with greater perceived boundary “salience”, measured by agreement across a separate group of individuals that a boundary occurred (Ben-Yakov & Henson, 2018). In addition to eliciting univariate responses, event boundaries have also been associated with multi-voxel pattern shifts within cortical areas during movie-viewing. Studies have reported that moments of transitions between neural states (i.e., shifts between stable multi-voxel patterns) in high-level cortical areas within the default mode network (DMN) significantly coincide with the moments identified as event boundaries by independent human raters, supporting the idea of *neural* event segmentation (Baldassano et al., 2017; Geerligs et al., 2021, 2022). The same studies also found that multi-voxel pattern shifts within the DMN overlapped with those in low-level cortical areas (e.g., the visual cortex) at event boundaries. This suggests a hierarchical structure in neural event segmentation, akin to the hierarchically structured coarse-grained and fine-grained event boundaries observed in earlier behavioral research (Zacks, Tversky, et al., 2001). However, despite the natural variation in the strength of event boundaries that individuals experience, such studies have tended to treat all human-identified event boundaries within a given dataset as equivalent. No studies to date have examined the relationship between event boundary *strength* and the magnitude of concomitant multi-voxel *pattern shifts* in cortical areas engaged by naturalistic movie-viewing.

Behaviorally, the probability of a person’s event segmentation in a narrative increases when the number of changes in situational features increases at that moment (Zacks, Speer, et al., 2009), which should correspond to perception of stronger event boundaries. Similarly, does neural event segmentation, indicated by transient changes in multi-voxel patterns, scale in a *graded* manner with event boundary strength in cortical areas? In contrast, a cortical region may merely detect the presence of an event boundary, with no modulation by boundary strength (i.e., a *binary* response profile). Importantly, even if neural responses (either univariate or pattern shift) at event boundaries are found to be graded at the group level, the question still remains whether they are graded at the individual subject level. This is because perceived event boundary strength could plausibly be correlated with the likelihood of boundary detection in a given brain area. If the brain area’s true response profile is binary, then averaging across subjects would yield an apparently graded response profile. That is, weak boundaries would produce the lowest score (e.g., detected by 20% of subjects), strong boundaries would produce the highest score (e.g., detected by 100% of subjects), and moderate boundaries would fall in between. Thus, analyses at the individual subject level are required to differentiate between graded and binary response profiles. Revealing these response properties of cortical areas may raise questions for models of event segmentation and event memory: if continuous experience is theorized to be segmented into discrete chunks, what purpose would graded cortical responses at boundaries serve?

In the current study, we first examined whether cortical regions across the brain showed a graded response profile in their event boundary-triggered pattern shifts, related to perceived boundary strength, during movie-viewing. We leveraged a publicly available naturalistic movie-viewing fMRI dataset (Lee & Chen, 2022). Event boundary strength was determined by agreement across an independent group of observers (following Ben-Yakov & Henson, 2018) and verified via direct judgment by another group. We organized our regions of interest according to a gradient of hierarchical timescales following the approach of prior studies (Baldassano et al., 2017; Geerligs et al., 2022). This was grounded in the theoretical claim that behavioral event segmentation concurrently occurs at multiple timescales (see Zacks et al., 2007). At multiple levels of the cortical hierarchy, we tracked changes in multi-voxel patterns at human-identified event boundaries grouped into three strength categories: weak, moderate, and strong. To assess whether each brain region’s activity was more compatible with a graded or a binary response profile, we analyzed the distributions of weak, moderate, and strong event boundary-triggered pattern shifts at the level of individual subjects. Next, motivated by prior findings that revealed a hierarchical structure in event boundary-related neural state transitions across the cortex (Baldassano et al., 2017; Geerligs et al., 2022), we tested whether the alignment of multi-voxel pattern transitions across cortical hierarchical levels was related to event boundary strength. Finally, we investigated whether the hippocampal response was greater at event boundaries when event boundary-triggered pattern shifts were more aligned across the cortical hierarchy.

## Methods

### Event segmentation data

Twenty-eight human observers performed an event boundary judgment task for the *Filmfest* movies. Fourteen undergraduate student participants (12 female, ages 18–22, mean age 19.6) were recruited from the Johns Hopkins SONA system and received credit for their participation. Additionally, 14 paid workers were recruited. We collected the data following a protocol approved by the Institutional Review Board of Johns Hopkins University. In the task, human observers were asked to identify boundaries between events with one-tenth-second resolution after a single complete viewing of the movie. In other words, event segmentation was performed retrospectively rather than through on-going segmentation during movie-viewing. Observers were instructed to place an event boundary whenever they perceived any meaningful change in scenes, such as shifts in topic, location, or characters (Newtson, 1973). The number of segments that observers were allowed to make varied for each movie, determined by a formula (the smallest integer greater than or equal to the movie duration in minutes multiplied by 3, plus or minus of that integer). For example, if a movie’s duration was 2 minutes 18 seconds, we asked them to segment 6 to 12 chunks. We also emphasized that they should use consistent decision criteria while making a judgment. To encourage precise judgment, they were allowed to watch a movie as many times as needed during the event segmentation task. We asked them to finish as many movies as possible within an allotted time frame. Event segmentation data were collected from 15-16 human observers for each movie following the sample size (*N* = 16) from Ben-Yakov and Henson (2018).

### Agreement-based event boundary strength

Event boundaries were identified at three levels of strength based on agreement across human observers (*weak*, *moderate*, and *strong* event boundaries). The more people placed an event boundary at a given time point in a movie, the stronger that event boundary was thought to be perceived. We first obtained a time series of event boundary agreement using methods from recent research (Michelmann et al., 2021). For each movie, each human observer’s boundary responses were entered into an array of time points, in which ‘1’ indicated the time point of an event boundary placed by that person. An inclusion window of 1s was used on each side of the actual event boundary (i.e., any time point within one second of an actual event boundary was also assigned a value of ‘1’). All other time points were assigned values of ‘0’. The arrays were averaged across people to obtain a time series of boundary agreement. The averaged time series was then smoothed using a Gaussian-weighted moving average filter with a 2-s window, and the time series data were concatenated across movies.

Event boundaries were identified by thresholding the concatenated, smoothed agreement data at the 65th percentile. After thresholding, the identified event boundaries were sorted by the degree of agreement, and divided into three bins (i.e., three categories of event boundary strength; **Figure 1**). Following Ben-Yakov and Henson (2018), pairs of event boundaries with distance closer than 6s were excluded from following fMRI analyses to avoid potential autocorrelation issues. In the fMRI data, movies were played consecutively. However, human observers performed event segmentation for each individual movie separately, resulting in inconsistent boundary judgments at the movie title onset (i.e., many subjects considered this the “start” of the movie and did not label a boundary there). Thus, we excluded event boundaries at movie onsets from fMRI data analyses. In addition, the last event boundary from each of the two movie-viewing runs was discarded due to the limited amount of brain data after that boundary. The times of event boundaries were shifted by 3 TRs to account for the hemodynamic response during fMRI data analyses.

**Figure 1.**
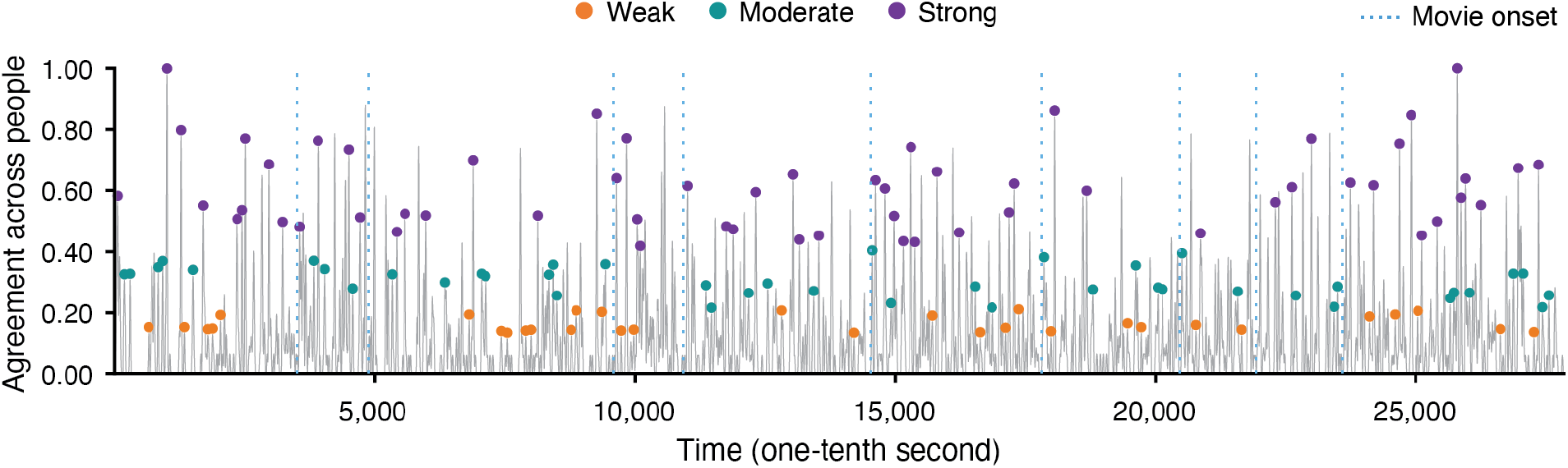
Three categories of event boundary strength. The gray line depicts the time series of the proportion of people agreeing on whether an event boundary occurred at each moment. The strength categories were based on the level of agreement across observers. The time point 0 corresponds to the onset of the first movie title, and subsequent movie title onsets are indicated by blue vertical dashed lines. Pairs of time points less than 6s apart were dropped. The time points of the 131 remaining event boundaries, analyzed in following fMRI analyses, are illustrated by colored dots (orange: weak, green: moderate, violet: strong).

### Event boundary strength rating

Four paid workers were provided with the event boundaries which were identified by the independent group of subjects described above. They then rated how strong each of the event boundaries was on a five-point scale (1: very weak, 5: very strong). After watching each movie, they examined each event boundary in the movie and judged its strength. This was done as a sanity check to verify that event boundary agreement corresponds to perceived boundary strength. Each rater’s responses were normalized, and then the normalized data were averaged across the raters. We computed a Spearman’s rank correlation between the group-average normalized rating and the event boundary agreement scores.

### Naturalistic movie-viewing fMRI dataset

We analyzed a publicly available fMRI dataset (*Filmfest*, Lee & Chen, 2022) in which participants watched a series of 10 short audiovisual movies with varying narrative content, plots, and emotions (3 cartoons and 7 live-action movies; movie duration range 2.15–7.75 minutes), in two scanning runs of five movies each, in fixed order. Further details about imaging data acquisition and parameters can be found in the original paper. The dataset was preprocessed using FSL (http://fsl.fmrib.ox.ac. uk/fsl) including motion correction, high-pass filtering (140s cutoff), spatial smoothing (4 mm FWHM Gaussian kernel) and registration to MNI standard space. For analyses, we resampled functional images to 3 mm isotropic voxels. Voxel time series were normalized using z-score within each scanning run. We discarded the data from four subjects due to excessive head motion (greater than 3 mm absolute displacement from the middle volume of the run). The dataset includes a recall session, but the current study used only the movie-viewing runs, and head motion was considered for these runs only. Data from 17 subjects were retained.

### Temporal receptive window (TRW) localizer and cortical hierarchy

#### TRW map

Four levels of the cortical hierarchy of processing timescales were defined by calculating the temporal receptive window (TRW) at the voxel level (Hasson et al., 2008; Lerner et al., 2011). The TRW refers to the duration preceding the present moment that influences brain responses, measuring the time over which information is accumulated in a cortical area. For example, if a brain area’s processing timescale is at the scale of a few seconds, its response at the present moment should not be affected by changing what information was presented a few seconds ago. TRWs are typically mapped across the brain by temporally scrambling a continuous natural stimulus at different levels of coarseness. The fMRI data used in the current TRW localizer analysis (*N* = 12) were taken from a prior study (Chen et al., 2016) in which an audiovisual movie stimulus (“Dog Day Afternoon”) was presented at three levels of temporal scrambling: (1) no-scramble (i.e., intact movie), (2) fine-scramble (temporal scrambling of 0.5–1.6-s segments), and (3) coarse-scramble (temporal scrambling of 7.1–22.3-s segments). We assessed each voxel’s across-subject response reliability (i.e., inter-subject correlation, ISC) by calculating correlations of voxel timecourses across brains within scrambling condition. Voxels were labeled as short-timescale (short-TRW) if they had above-threshold ISC (*r* > .2, as in Chen et al., 2016) in all three scrambling conditions; as medium-timescale (medium-TRW) if above-threshold ISC was observed both in the coarse-scramble and the no-scramble conditions, but not in the fine-scramble condition; and as long-timescale (long-TRW) if they had above-threshold ISC only in the no-scramble condition. In this way, three initial levels of cortical hierarchy were defined.

#### Parcel assignment

Short-, medium-, and long-timescale voxels were separated into corresponding cortical parcels from 17 brain networks in a 400-parcel atlas derived from resting-state functional connectivity data (Schaefer et al., 2018). First, parcels were rejected if fewer than 50% of their voxels overlapped with the TRW map (any combination of short, medium, and long timescale voxels). For each remaining parcel, a timescale label of short, medium, or long was assigned based on which described the greatest number of voxels for that parcel.

Short-timescale cortical parcels were further divided into two sub-groups based on whether they belonged to a modality-specific area (early visual cortex or auditory cortex) or not. Parcels were defined as being in early visual cortex if more than 50% of the voxels within a parcel overlapped with areas V1–V4 in a probabilistic atlas (L. Wang et al., 2015; threshold = .25). Auditory cortex parcels were hand-selected based on anatomical correspondence to Heschl’s gyrus. These parcels in early sensory processing areas (visual and auditory) were assigned to level 1 of the cortical hierarchy. The rest of the short-timescale parcels were defined as level 2 of the cortical hierarchy. This resulted in four levels of cortical hierarchy (**Figure 2A**; level 1: visual or auditory processing areas with a short timescale, level 2: non-sensory-specific areas with a short timescale, level 3: cortical areas with a medium timescale, level 4: cortical regions with a long timescale).

**Figure 2.**
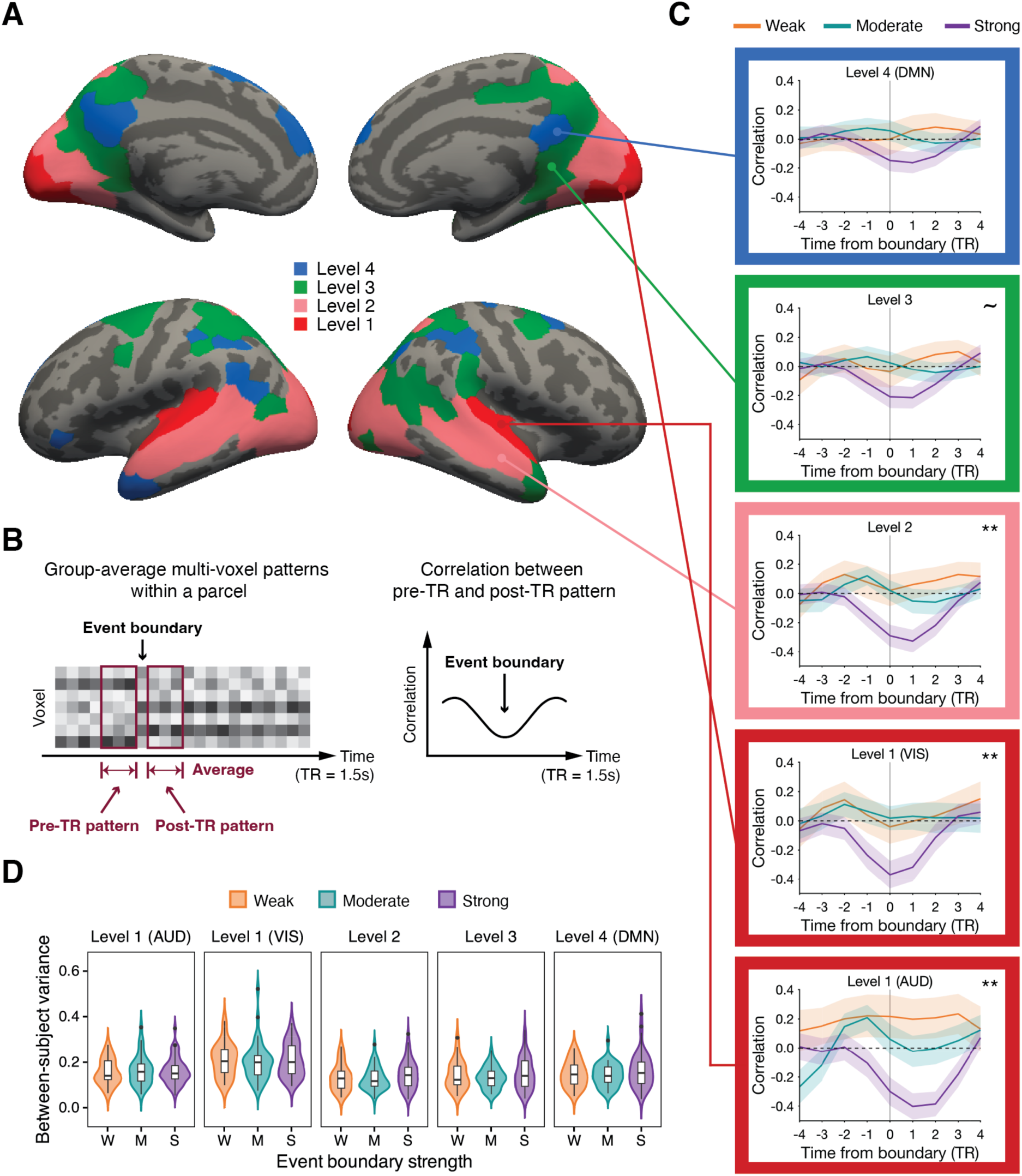
Event boundary-locked changes in multi-voxel patterns along the cortical hierarchy. (A) Cortical hierarchy of processing timescales. (B) For each time point (TR) in the group-average data, we computed a pre-TR and post-TR pattern (left). The correlation between the pre-TR and post-TR patterns was calculated for every TR (right), normalized across the entire scan. (C) Changes in multi-voxel patterns around event boundaries for different strengths (gray vertical line: event boundary, AUD: auditory processing areas, VIS: visual processing areas, DMN: the default mode network, shaded area: ± SEM across event boundaries, ∼ *p* < .1, ** *p* < .01 main effect of boundary strength). Note that correlation time series was normalized (see Methods). (D) For each event boundary, the between-subject variance in cross-boundary correlation values was computed. The distribution of between-subject variance is illustrated (black horizontal bar: median, black dot: outlier). We performed a Bayesian one-way ANOVA, which revealed evidence in favor of the null model: no differences in between-subject variance across event boundary strength groups at any cortical hierarchical level.

The long-timescale cortical areas at level 4 (indicated in blue in **Figure 2A**) broadly overlapped with core regions within the DMN, including the bilateral posterior medial cortex, bilateral medial prefrontal cortex, left angular gyrus, and left temporal pole. 68.9% of the voxels (55.6% of the parcels) at level 4 belonged to the DMN, based on the labels from the 17-network atlas. A prior study (Chen et al., 2016), which provided the TRW localizer data, also reported that these areas in the original TRW map coincide with the main regions in the DMN.

### Cortical pattern shift time series

For each parcel at every cortical hierarchical level we defined, we tracked shifts in multi-voxel patterns at each time point (repetition time, TR) of the fMRI data.

#### Group-average analyses

At each TR, we calculated both pre– and post-TR patterns by averaging the multi-voxel patterns across 3 TRs separately for the periods immediately before and after a given TR (**Figure 2B**, left). Then, a correlation between the pre– and post-TR patterns was computed for that TR. By repeating these procedures at each TR in both movie-viewing runs, we generated a correlation time series of cortical multi-voxel pattern shift. For each parcel, the correlation time series from each scanning run was concatenated, and then the concatenated time series was normalized (z-scored) across the two movie-viewing runs. Next, the normalized data were averaged across parcels at each cortical hierarchical level. In other words, we combined z-scores from different parcels at the same cortical hierarchical level. We refer to this time series of combined z-scores as a parcel-average correlation time series.

The parcel-average correlation values for the 4 TRs preceding and following each event boundary are illustrated in **Figure 2C** for visualization purposes. In this study, a “cortical boundary shift” denotes a change in multi-voxel patterns at a human-judged event boundary. To measure the magnitude of cortical boundary shifts, we defined “cross-boundary correlation” as the mean correlation averaged across –1 to 1 TRs from each event boundary at each cortical hierarchical level. These cross-boundary correlation values were computed using the parcel-average correlation time series. We found that the cross-boundary correlation data in the early auditory cortex and the regions at level 3 deviated from the normality assumption from the Shapiro-Wilk test. Thus, instead of conducting one-way ANOVA, we performed a non-parametric Kruskal-Wallis test on the cross-boundary correlation data separately for each cortical hierarchical level, using boundary strength categories: weak, moderate, and strong. Subsequently, a Dunn’s test was carried out for pairwise comparison.

Furthermore, we calculated a Spearman’s rank correlation between the two continuous variables of cross-boundary correlation and across-observer event boundary agreement at each level of the cortical hierarchy. In this analysis, we used the continuous value of across-observer event boundary agreement, rather than binning it into the three boundary strength categories (weak, moderate, and strong).

#### Individual-subject analyses

For individual subject analyses, we obtained a cortical pattern shift time series using each subject’s multi-voxel patterns data, instead of the group-average data. Analysis procedures were otherwise identical to those described above (see *Group-average analyses*). For each subject, a cross-boundary correlation value was calculated for each event boundary. At each event boundary, the between-subject variance in cross-boundary correlation was computed (**Figure 2D**). To examine whether the between-subject variance varies according to boundary strength, a Bayesian one-way ANOVA was performed using JASP (Version 0.18.3; JASP Team, 2024). Two models were tested: the null model and the event boundary strength model. Since we had no prior information, the prior for each model was initially set to a default value of 0.5, assuming that each model was equally likely at the beginning.

We performed Hartigan’s dip test (Hartigan & Hartigan, 1985) on the distribution of each subject’s cross-boundary correlation data separately for each event boundary strength category at each cortical hierarchical level (**Figure 3**). This was done to examine whether the distributions were unimodal or not, which was necessary for testing the idea of graded vs. binary response profiles in the cortical boundary shift. We ran a one-way repeated measures ANOVA on individual-subject cross-boundary correlation data to test the differences in the means among boundary strength categories.

**Figure 3.**
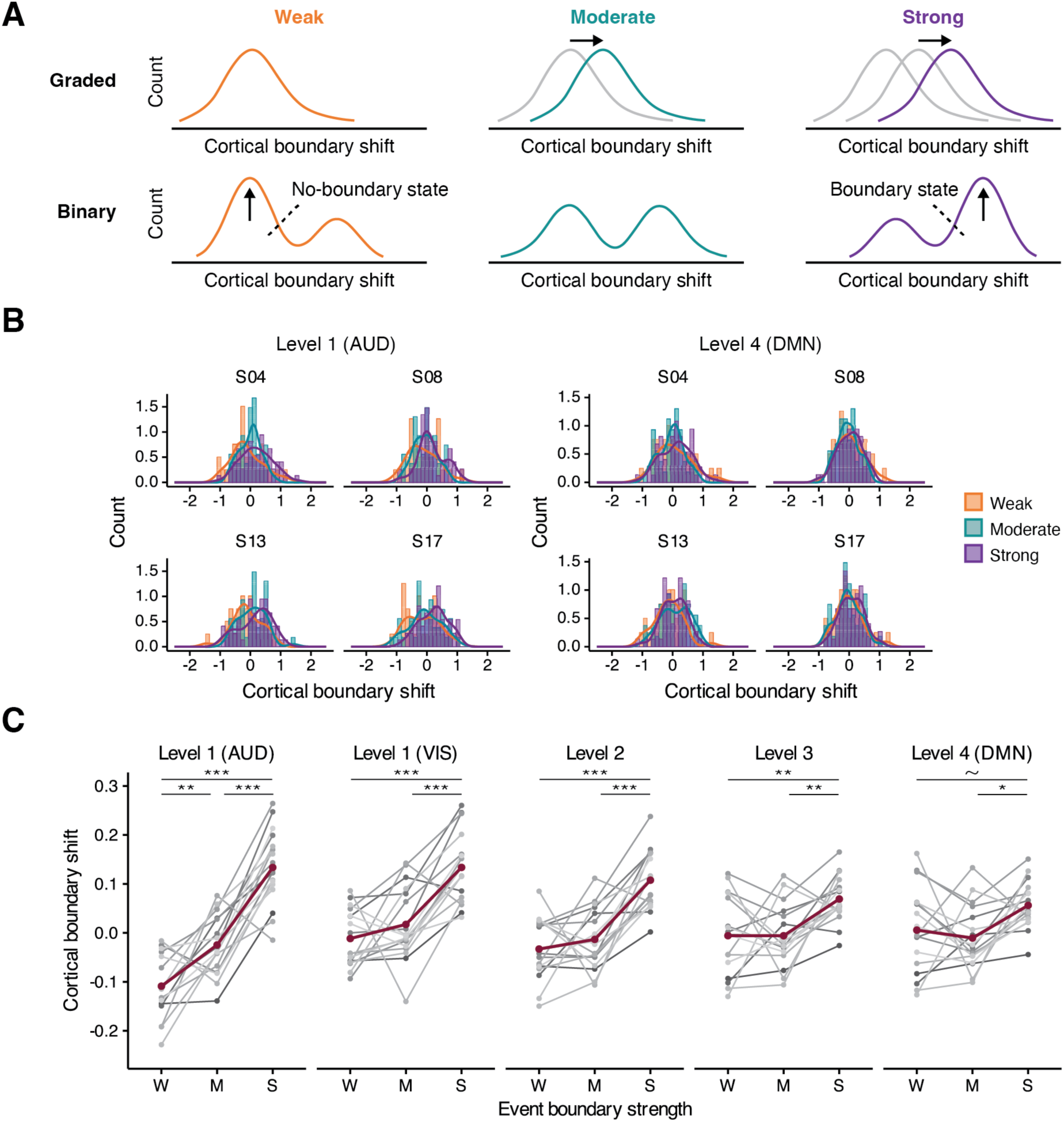
Graded vs. binary response profiles. (A) Predicted distributions of cortical boundary shift values for different boundary strength categories for the graded and binary response profiles. A brain region with a binary response profile should exhibit a bimodal distribution shape, suggesting the presence of two distinct states: no-boundary and boundary states. (B) The distribution data from four selected subjects in auditory processing areas (AUD) at level 1 (left) and the default mode network (DMN) at level 4 (right). No clear evidence for bimodality was observed. In auditory processing areas, distributions match the predictions for a graded response profile. (C) The means of the cortical pattern shift values in each event boundary strength category. Gray shades depict individuals’ data; red lines illustrate group means. ∼ *p* < .1, * *p* < .05, ** *p* < .01, *** *p* < .001 post-hoc pairwise comparison (Bonferroni correction). W: weak; M: moderate; S: strong

#### Non-boundary response analysis

In order to investigate if the distribution of individuals’ cross-boundary correlation shows bimodality (i.e., a binary response profile) during non-boundary periods, we also sampled a total of 44 time points (22 for each movie-viewing run) approximately matching the average number of event boundaries from the boundary strength categories. The sampling was pseudo-randomly performed, excluding time points during non-movie periods (e.g., movie title scenes), and dropping any new time point closer than 6s to a previously sampled time point to prevent potential autocorrelation issues for adjacent time points. Additionally, time points during the last 10s of each movie-viewing run were excluded to ensure in-movie time points were sampled. After sampling these non-boundary time points, Hartigan’s dip test was performed again on the distribution of each subject’s cross-boundary correlation data during these non-boundary periods.

### Cortical alignment score of pattern shift

We designed methods for scoring the degree of alignment of multi-voxel pattern shifts across cortical hierarchical levels at human-identified event boundaries. A higher score indicates that the multi-voxel pattern shifts were aligned across more cortical hierarchical levels at a given event boundary (**Figure 4A**, top). The presence of a shift was defined as a cross-boundary correlation value smaller than 0. Three types of cortical alignment scores (*strict nesting*, *nesting*, and *summation*) were computed based on whether a bottom-up nested structure in cortical boundary shifts was present, with varying degrees of strictness. For *nesting* and *summation*, modality-specific areas were not included in the scoring process in order to make the measures more lenient. The results from the *nesting* scoring method are reported in **Figure 4** and those from the *strict nesting* or *summation* can be found in **Supplementary Figure 7**.

**Figure 4.**
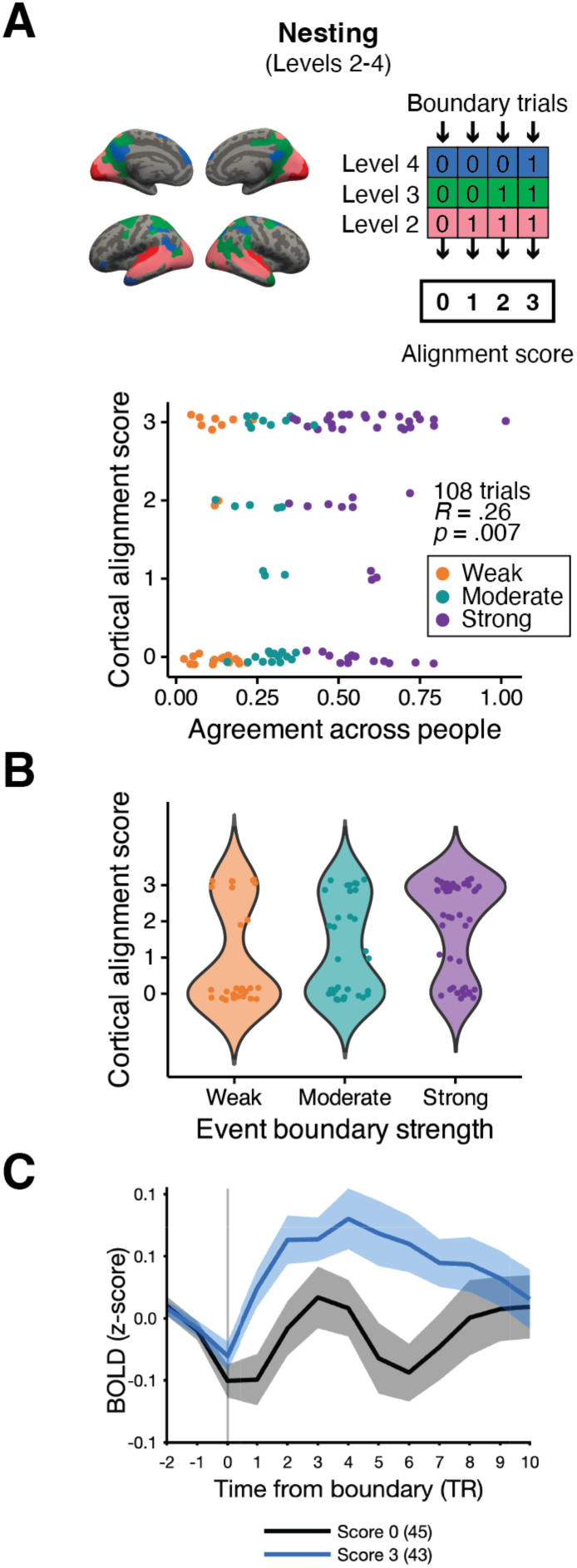
Alignment of multi-voxel pattern shifts across the cortical hierarchy and event boundary strength. Event boundaries were assigned a score representing the degree of alignment of pattern shifts across cortical hierarchical levels (the nesting method). (A) We observed a significant positive relationship between the degree of cortical alignment and across-observer boundary agreement. Individual dots in the scatter plot illustrate event boundary trials. (B) The same data is presented in a violin plot to visualize the distribution. (C) Each line illustrates hippocampal activity at event boundaries with different alignment scores (blue solid line: the highest possible alignment score, i.e., score 3 for the nesting method; black solid line: score 0; shaded area: ± SEM across subjects). In the legend, the number in a parenthesis shows the number of event boundary trials associated with that score.

We first constructed a cortical level-by-boundary data matrix in which a value of ‘1’ was assigned if a below-zero cross-boundary correlation value was found at a specific cortical hierarchical level at a given human-identified event boundary. Otherwise, each cell in the data matrix contained ‘0’. The calculation of cortical alignment score was based on the group-average fMRI data using the values in the data matrix for each event boundary. In *strict nesting*, we evaluated the presence of pattern shifts at all four cortical hierarchical levels in a bottom-up nested manner (score range 0–4). For example, score 4 indicates the presence of pattern shifts at all cortical hierarchical levels we defined. In *nesting*, the procedures remained the same, but only levels 2–4 were assessed (score range 0–3); a cortical boundary shift in modality-specific areas was not taken into account. In *summation*, the bottom-up nested structure in pattern shifts was not considered (score range 0–3). For instance, a score of 2 was assigned to an event boundary if there were pattern shifts at level 2 and level 4, or at level 3 and level 4. Finally, a Spearman’s rank correlation was computed between each type of cortical alignment score and the event boundary agreement data (**Figure 4A**, bottom, for *nesting*). We also plotted the distribution of alignment scores for each event boundary strength category (**Figure 4B**, for *nesting*).

### Hippocampal response at event boundary

To explore hippocampal activity at event boundaries, we anatomically defined bilateral hippocampus using the Harvard-Oxford atlas (Desikan et al., 2006). Event boundaries were grouped by cortical alignment scores, ranging from 0 to 4 for *strict nesting* or 0 to 3 for either *nesting* or *summation*, instead of event boundary strength categories. The analysis was performed at the individual level. We extracted a time series of hippocampal activity for each subject and each event boundary, covering the period from 2 TRs before to 10 TRs after the event boundary. The baseline activity was calculated by averaging the data for the 2 TRs preceding the event boundary. This baseline activity was next subtracted from the data at each TR following the prior study which provided the current naturalistic movie-viewing dataset (Lee & Chen, 2022). Hippocampal activity from 0 to 4 TRs after an event boundary was averaged, and then averaged again across event boundaries with the same alignment score. A one-way repeated measures ANOVA or a paired *t*-test was conducted to test for differences in the means among all cortical alignment scores or between a pair of scores (**Figure 4C**, for *nesting*; **Supplementary Figure 7**, for all score types).

The same hippocampal response analysis was performed for event boundaries in the three strength categories, as in a prior study (Ben-Yakov & Henson, 2018), to examine whether we could replicate the boundary strength modulation in our dataset. For this and all other such analyses in the current study, we used a 2-TR baseline preceding event boundaries (Lee & Chen, 2022) because we felt that the movies in our dataset were fast-paced compared to those used in other studies. Since earlier research used a baseline of 10 TRs (Baldassano et al., 2017), we additionally calculated hippocampal responses using a 10-TR baseline in order to facilitate comparison between the studies.

## Results

### Event boundaries grouped by strength

Event boundary strength was determined by the degree of agreement across individuals (an independent group from those who participated in the fMRI study), following the approach used in prior research (Ben-Yakov & Henson, 2018). Using peak boundary detection methods from Michelmann and colleagues (2021), a total of 314 boundaries were initially identified for the *Filmfest* movies (see *Agreement-based event boundary strength* in Methods). These event boundaries were divided into three categories of strength (see **Supplementary Table 1**). For example, at one point in the movie “Catch Me If You Can”, the video abruptly shifts from a gameshow studio stage to a rainy outdoor night in Marseille; almost all observers (98%) labeled this as an event boundary. In prior research, event boundaries were included if at least five out of 16 people (31.3%) agreed (Ben-Yakov & Henson, 2018; Geerligs et al., 2022). In this study, however, we sought to capture a wider range of strength, and thus incorporated a wider range of event boundary agreement levels (at least 11% from the smoothed event boundary agreement data; the raw numbers vary because there was a different number of raters for each movie). We removed boundaries which might introduce unwanted noise: all boundaries which were less than 6s from another boundary (both removed, as in Ben-Yakov & Henson, 2018), boundaries at movie title onsets, and the last boundary from each scanning run (see *Agreement-based event boundary strength* in Methods for details). Therefore, a total of 131 event boundaries were included in subsequent fMRI analyses (31 *weak* event boundaries; 42 *moderate* event boundaries; 58 *strong* event boundaries; **Figure 1**; see **Supplementary Table 2** for the agreement range).

To confirm that perceived boundary strength was related to agreement, four paid workers rated boundary strength on a five-point scale (1: very weak, 5: very strong). We found a strong positive relationship between event boundary consensus and boundary strength ratings of the 131 event boundaries (*r*(129) = .64, *p* < .001).

### Larger multi-voxel pattern transitions were observed at stronger event boundaries

Prior studies reported that human-labeled event boundaries significantly coincided with the moments of neural state transitions in the DMN (Baldassano et al., 2017; Geerligs et al., 2021). In these studies, neural state transitions were identified using computational algorithms which searched for transient changes in multi-voxel cortical patterns, and then the overall temporal match between human-labeled and algorithm-labeled boundary lists was statistically evaluated. In the current study, we instead anchored on human-identified event boundaries and examined multi-voxel pattern changes at these moments. We examined pattern change dynamics at event boundaries in brain parcels at four levels spanning the cortical hierarchy of timescales, defined by an independent localizer (see **Figure 2A** for the cortical hierarchy; see *Temporal receptive window localizer and cortical hierarchy* in Methods). Specifically, we asked whether the magnitude of cortical pattern shift was greater for event boundaries which were perceived to be stronger. A time series of multi-voxel pattern shifts was obtained at each cortical hierarchical level using the group-average data (see *Cortical pattern shift time series* in Methods for details). This approach minimizes idiosyncratic noise and maximizes stimulus-driven signals, since all subjects viewed the exact same movies (Baldassano et al., 2017; Geerligs et al., 2021). Each element in the time series was a correlation coefficient between a pair of multi-voxel patterns, one window immediately prior and one immediately following that time point (**Figure 2B**). These were averaged across parcels to produce one time series for each of the four hierarchical levels.

**Figure 2C** illustrates the average time series of multi-voxel pattern shifts around event boundaries. Visual inspection suggests that the multi-voxel pattern transiently changes (i.e., decreased correlation) at event boundaries, then stabilizes (i.e., increased correlation) after the boundary, consistent with findings from prior research (Baldassano et al., 2017; Geerligs et al., 2021). To evaluate whether cortical boundary pattern shifts differ depending on boundary strength, a non-parametric Kruskal-Wallis test was performed at each cortical hierarchical level separately (see *Cortical pattern shift time series* in Methods). Here, we refer to the correlation coefficient averaged across –1 to 1 TRs from an event boundary as a “cross-boundary correlation”. Note that a smaller or more negative value of cross-boundary correlation indicates a greater magnitude multi-voxel pattern shift (i.e., cortical boundary shift). We found a significant main effect of boundary strength on cortical boundary shift in short-timescale cortical regions (ᵡ^2^(2, *N* = 131) = 11.65, *p* = .003, η^2^ = .08 in auditory processing areas at level 1; ᵡ^2^(2, *N* = 131) = 10.06, *p* = .007, η^2^= .06 in visual processing areas at level 1; ᵡ^2^(2, *N* = 131) = 10.37, *p* = .006, η^2^ = .07 in the cortical regions at level 2). We observed a trending main effect of boundary strength in the cortical areas at level 3 (ᵡ^2^(2, *N* = 131) = 5.76, *p* = .056, η^2^ = .03), but the main effect was not significant at level 4 (ᵡ^2^(2, *N* = 131) = 4.29, *p* = .117, η^2^ = .02).

In auditory processing areas and the short-timescale regions at level 2, the means of cross-boundary correlation, averaged across the event boundary trials within a strength category, were in a graded order, with a significant difference between strong and weak, and between strong and moderate (auditory processing areas: *M* = –0.27, *SD* = 0.65 for strong; *M* = 0.08, *SD* = 0.59 for moderate; *M* = 0.21, *SD* = 0.75 for weak; level 2: *M* = –0.26, *SD* = 0.53 for strong; *M* = 0.03, *SD* = 0.41 for moderate; *M* = 0.05, *SD* = 0.54 for weak). The statistical results from the post-hoc pairwise comparison are presented in **Table 1**. A similar pattern was found in visual processing areas, but the means were not in a perfectly graded order (visual processing areas: *M* = –0.31, *SD* = 0.66 for strong; *M* = 0.04, *SD* = 0.48 for moderate; *M* = 0.00, *SD* = 0.58 for weak). No significant difference between weak and moderate event boundaries was found in any short-timescale regions. While level 3 and level 4 regions did not show main effects of boundary strength, numerically the means appeared in graded order at level 4 (level 3: *M* = –0.18, *SD* = 0.53 for strong; *M* = 0.02, *SD* = 0.44 for moderate; *M* = –0.01, *SD* = 0.55 for weak; level 4: *M* = –0.13, *SD* = 0.51 for strong; *M* = 0.05, *SD* = 0.41 for moderate; *M* = 0.02, *SD* = 0.58 for weak).

**Table 1.** Post-hoc pairwise comparison (Dunn’s test) results in the group-level analysis. Bonferroni correction was applied. The corrected *p*-value is the initial *p*-value multiplied by the number of comparisons. Note that no significant main effect was observed at level 4 (DMN) nor at level 3.

| Pairwise comparison |  | Z | p (corrected) |
| --- | --- | --- | --- |
| Level 1 (AUD) | Weak vs. Moderate | -0.63 | 1 |
|  | Weak vs. Strong | -3.04 | .007 |
|  | Moderate vs. Strong | -2.60 | .028 |
| Level 1 (VIS) | Weak vs. Moderate | 0.12 | 1 |
|  | Weak vs. Strong | -2.43 | .045 |
|  | Moderate vs. Strong | -2.81 | .015 |
| Level 2 | Weak vs. Moderate | -0.24 | 1 |
|  | Weak vs. Strong | -2.68 | .022 |
|  | Moderate vs. Strong | -2.67 | .023 |

In the subsequent correlation analysis, we further investigated if there is a significant relationship between the magnitude of cortical boundary shift and the continuous across-observer event boundary agreement (i.e., strength) at each level of the cortical hierarchy (**Supplementary Figure 1**). This extends our group-level analysis shown in Figure 2C by using a continuous rather than a categorical variable for boundary strength. Note that a negative relationship is expected in this analysis due to the fact that a smaller value of cross-boundary correlation indicates a larger transition in multi-voxel patterns, which is thought to be associated with a stronger event boundary. Consistent with the results from the preceding analysis, the relationship was the strongest in short-timescale, auditory processing areas at level 1 (*r*(129) = –.36, *p* < .001), whereas the relationship became weaker at level 4 (*r*(129) = –.16, *p* = .070).

In summary, we observed that event boundary-related pattern shifts in short-timescale cortical areas (early auditory cortex and early visual cortex) exhibited a response at event boundaries modulated by boundary strength. This was pronounced for the discrimination between strong and weak, and between strong and moderate event boundaries. The differences between strength categories were not significant in higher-level cortical areas at level 3 and level 4. The results from the subsequent correlation analysis using the continuous across-observer event boundary agreement values further confirmed that the association between cortical boundary shift and event boundary agreement progressively decreased in higher-level cortical areas with longer processing timescales.

### Graded vs. binary response profiles

#### Between-subject variance in cortical boundary responses

In the preceding analysis, we averaged data across subjects before calculating pattern shift timecourses. However, it leaves open the possibility that, for regions which appeared to have a graded response profile, the gradation was due to increasing likelihood of “detection” rather than to a gradually increasing neural response with increasing boundary strength. That is, each individual subject’s neural responses may have been “binary”, with a pattern shift occurring when a boundary was “detected” but not scaling with boundary strength; the higher the likelihood that any given subject would detect the boundary, the greater the average response magnitude would be. We sought to distinguish between these two alternatives (graded vs. binary response profiles) by examining the variance of the cortical pattern shift across subjects within each boundary strength category (weak, moderate, strong). We reasoned that, under the “binary” account, between-subject variability should be the highest for moderate boundaries, as subjects would be maximally divided across “detected” and “undetected”. We expected a “no-boundary state” in most people for the weak boundary category, resulting in relatively low between-subject variance, similar to the strong boundary category.

For this analysis, we obtained a time series of cortical pattern shift using each subject’s own data (see *Individual-subject analyses* in Methods). For every event boundary trial, we calculated the variance in cortical boundary shift across subjects. The key comparison was whether the distribution of between-subject variance from the moderate boundary category was significantly different from that of strong and weak categories. Figure 2D illustrates the distribution of between-subject variance in each boundary strength category. To investigate whether there were differences in between-subject variance of cortical boundary shift among boundary strength groups, we performed a Bayesian one-way ANOVA for each cortical hierarchical level (see *Individual-subject analyses* in Methods). However, we observed strong evidence in favor of the null model hypothesis (i.e., graded response) in auditory processing areas (*BF*_01_ = 11.89; *M* = –0.003, *SD* = 0.007, 95% CI = [-0.018, 0.011] for weak; *M* = 0.002, *SD* = 0.007, 95% CI = [-0.012, 0.016] for moderate; *M* = 7.60 × 10^-4^, *SD* = 0.006, 95% CI = [-0.012, 0.014] for strong), and visual processing areas (*BF*_01_ =12.14; *M* = 0.002, *SD* = 0.010, 95% CI = [-0.017, 0.021] for weak; *M* = –0.003, *SD* = 0.009, 95% CI = [-0.021, 0.014] for moderate; *M* = 0.001, *SD* = 0.008, 95% CI = [-0.016, 0.017] for strong) at level 1. For example, the between-subject variance data in auditory processing areas was 11.89 times more likely to occur under the null model than the event boundary strength model. Moderate evidence in favor of the null model was also observed in cortical areas at level 2 (*BF*_01_ = 4.57, *M* = –0.003, *SD* = 0.007, 95% CI = [-0.018, 0.011] for weak; *M* = –0.006, *SD* = 0.007, 95% CI = [-0.020, 0.007] for moderate; *M* = 0.009, *SD* = 0.006, 95% CI = [-0.004, 0.022] for strong), level 3 (*BF*_01_ = 5.79, *M* = 5.11 × 10^-4^, *SD* = 0.008, 95% CI = [-0.015, 0.016] for weak; *M* = –0.008, *SD* = 0.007, 95% CI = [-0.022, 0.006] for moderate; *M* = 0.007, *SD* = 0.007, 95% CI = [-0.007, 0.021] for strong) and level 4 (*BF*_01_ = 4.97, *M* = –0.003, *SD* = 0.008, 95% CI = [-0.019, 0.013] for weak; *M* = –0.007, *SD* = 0.007, 95% CI = [-0.022, 0.008] for moderate; *M* = 0.009, *SD* = 0.007, 95% CI = [-0.005, 0.023] for strong). Note that the means are posterior distribution means.

#### Distribution shape

We performed a second test of the graded vs. binary response profile accounts by examining the shapes of the distributions of cortical pattern shifts from the three boundary strength categories at the individual subject level. The binary response profile account predicts bimodality in the distribution of cortical boundary shift because the values should fall into two bins: event boundary detected, or no boundary detected (Figure 3A). In contrast, the graded response profile account predicts that the distributions should be unimodal, with the means arranged in order of event boundary strength (weak < moderate < strong). More negative cross-boundary correlation values indicate larger transitions at event boundaries; to visualize cortical boundary shifts in Figure 3, we flipped the sign of the cross-boundary correlation values for ease of interpretation, so that a larger value would correspond to a greater cortical boundary shift. The sign flipping did not affect any statistical tests.

**Figure 3B** illustrates the distribution of cortical boundary shift in auditory processing areas at level 1 and DMN regions at level 4 in the three boundary strength categories. Data from four subjects are presented for visualization purposes (see **Supplementary Figures 2-6** for the distributions from all subjects at each cortical hierarchical level). Using Hartigan’s dip test, we evaluated whether the distribution of each boundary strength category is unimodal or not, for each subject at each cortical hierarchical level. None of the data distributions were significantly different from the statistical null hypothesis of a unimodal distribution, thus providing no evidence for the binary response profile account. The same Hartigan’s dip test was performed on non-boundary trials, but the null hypothesis of unimodality was not rejected (see *Non-boundary response analysis* in Methods for details about the sampling of non-boundary trials). The gray line in **Supplementary Figures 2-6** indicates a density curve of the non-boundary data.

Next, we examined whether increased event boundary strength is associated with a progressive change in cortical boundary shift values, in each individual’s data. A one-way repeated measures ANOVA was run on the cortical boundary shift data, separately for each cortical hierarchical level. We observed a significant main effect of event boundary strength at all cortical hierarchical levels (*F*(2, 32) = 62.35, *p* < .001 in auditory processing areas; *F*(2, 32) = 24.74, *p* < .001 in visual processing areas; *F*(2, 32) = 29.28, *p* < .001 at level 2; *F*(2, 32) = 8.74, *p* < .001 at level 3; *F*(2, 32) = 5.09, *p* = .012 at level 4). The effect size decreased at higher levels of the cortical hierarchy, such as in DMN regions at level 4 (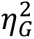 = .71 in auditory processing areas; 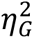 = .47 in visual processing areas; 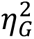 = .52 at level 2; 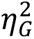 = .24 at level 3; 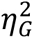 = .15 at level 4). Post-hoc pairwise comparison revealed significant differences between every pair of event boundary strength categories only in auditory processing areas at level 1, which have a short processing timescale; that is, there was a clear gradient of cortical boundary shift in auditory processing areas at level 1 (*M* = –0.11, *SD* = 0.06 for weak; *M* = –0.03, *SD* = 0.06, for moderate; *M* = 0.13, *SD* = 0.08 for strong). There was no difference between weak and moderate in other regions of the cortical hierarchy, including the visual cortex. In these regions, the effect of event boundary strength was primarily driven by the differences between either strong and moderate, or strong and weak event boundaries. The results from the post-hoc pairwise comparison can be found in **Table 2**. In Figure 3C, individual lines in shades of gray show each subject’s data at each cortical hierarchical level.

**Table 2.**
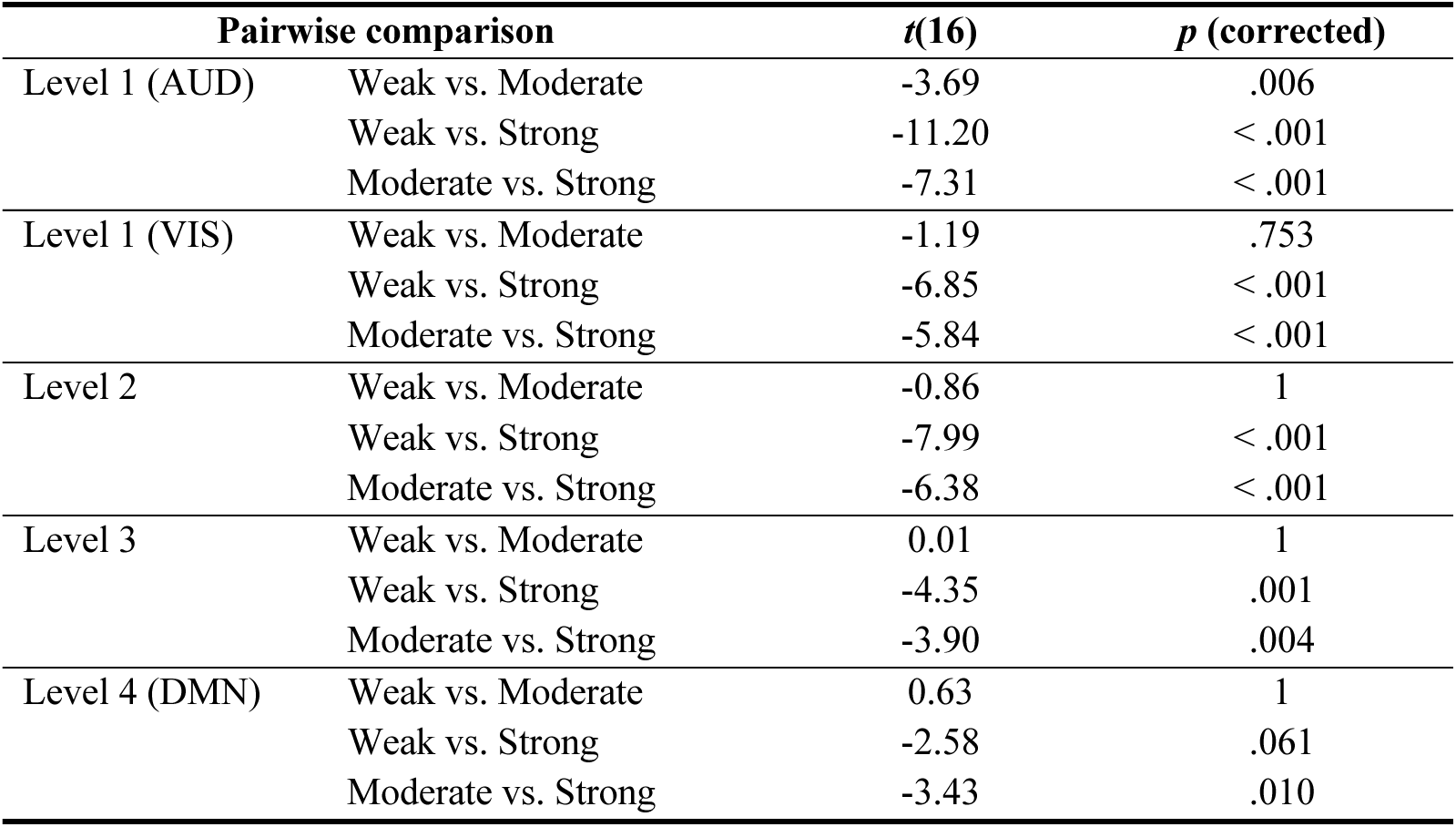
Post-hoc pairwise comparison (paired *t*-test) results in the individual-level analysis. Bonferroni correction was applied. The corrected *p*-value is the initial *p*-value multiplied by the number of comparisons.

In the above analyses, we tested whether between-subject variance in cortical boundary shift is greater for moderate event boundaries which we assumed would result in the greatest separation in boundary agreement across individuals under a binary response profile. There is a possibility that the true boundary strength category with the maximal separation could be something other than moderate event boundaries (e.g., a sub-group within the strong event boundary category). Nevertheless, we did not observe differences in between-subject variance among boundary strength categories at any cortical hierarchical level.

Taken together, our investigations using individual subject analyses provide support for a strongly graded response profile in auditory processing areas, which have a short timescale, consistent with the results from the group-level analyses. Event boundary strength also modulated pattern shifts in higher level cortical areas (levels 2, 3 and 4), with significant differentiation between strong and moderate boundary strength, but not between weak and moderate. No evidence was found supporting the binary response account at any level of the cortical hierarchy from the investigation of distribution shapes at the individual subject level; however, note that these were null results, not constituting strong evidence *against* the binary response account.

### The relationship between cortical alignment in pattern shifts and event boundary strength

In the above analyses, we tested whether brain areas discriminate event boundary strength within each cortical hierarchical level. Next, we sought to examine whether event boundary strength modulates how much boundary-related neural responses *align across different levels* of the cortical hierarchy of timescales. Specifically, we asked whether there was a positive association between 1) the extent of alignment across levels, in terms of multi-voxel pattern shifts, and 2) event boundary strength. In prior studies that investigated the relationship between nested cortical neural state transitions and behaviorally judged event boundaries, all human-identified event boundaries were treated equally (Baldassano et al., 2017; Geerligs et al., 2022). Here, however, an agreement value was associated with each human-identified event boundary. We asked whether an event boundary is perceived to be stronger when pattern shifts align more across cortical hierarchical levels at that boundary.

To address these questions, for each event boundary, we first defined a method of calculating cortical alignment (see *strict nesting*, *nesting*, and *summation* in *Cortical alignment score of pattern shift* in Methods). These calculations of alignment required, for a given cortical hierarchical level, the labeling of each time point as either “shift” or “no shift”. We labeled a time point as “shift” if the cross-boundary correlation was less than 0 (see *Cortical pattern shift time series* in Methods for the definition of “cross-boundary correlation”). The magnitude of cortical pattern shift was not considered during scoring procedures. For example, in the strict nesting method, an event boundary trial was classified as *not* exhibiting a bottom-up nested structure if there were pattern shifts at levels 2 and 3 but not at level 1. In the *nesting* method, a bottom-up nested structure was evaluated for levels 2 and higher of the cortical hierarchy, ignoring level 1. Nested structure was irrelevant in the summation method. Figure 4A shows the main results from the nesting method. Cortical alignment was significantly correlated with event boundary strength (*r*(106) = .26, *p* = .007). The results from the strict nesting and the summation methods were consistent with the nesting method (**Supplementary Figure 7**; *r*(101) = .26, *p* = .008 for strict nesting; *r*(129) = .23, *p* = .008 for summation). The distribution of cortical alignment scores from the nesting method is also illustrated in Figure 4B according to the three event boundary strength categories. Cortical alignment scores in the strong event boundary category were generally higher compared to either the weak or moderate category.

We noticed that only a small portion of event boundaries (17.6%; 23 out of 131) lacked a bottom-up nested structure within level 2 to level 4 of the cortical hierarchy and were excluded from our main nesting scoring procedures. Also, it was most common for cortical pattern shifts to be aligned across *all* cortical hierarchical levels, as opposed to being more sparsely aligned. For example, when using the strict nesting method, 32.8% of event boundaries were assigned a score of 4 (score 0: 19.1%; score 1: 15.3%, score 2: 3.1%, score 3: 8.4%). Those event boundaries with score 4 from the strict nesting method were identical to those with the highest score (i.e., score 3) in the nesting method, wherein the presence of cortical boundary shift in modality-specific, sensory processing areas (i.e., level 1) was not considered during the scoring procedures.

In summary, we found a significant positive relationship between the degree of alignment of multi-voxel pattern shifts across the cortical hierarchy and event boundary strength. In addition to cortical areas responding to boundary strength in a graded manner, the presence of multi-voxel pattern shifts at multiple cortical hierarchical levels is also associated with gradual changes in boundary strength. Both a graded response within a cortical region and the alignment of multi-voxel pattern shifts across distinct cortical areas with different processing timescales were correlates of event boundary strength. Because a small number of event boundaries lacked a bottom-up nested structure of multi-voxel pattern shifts, we could not further compare the results from the summation method to either the strict nesting or nesting methods.

### Hippocampal response at event boundaries with different cortical alignment scores

As Ben-Yakov and Henson (2018) reported that hippocampal responses during movie-viewing not only increased at event boundaries but were modulated by boundary salience, we examined hippocampal event boundary responses in our data. Continuing our investigations from the prior section, we asked: Are hippocampal responses at event boundaries related to the degree of alignment of multi-voxel pattern shifts across different levels of the cortical hierarchy? Instead of grouping event boundaries by boundary strength categories, we grouped them by cortical alignment scores (see *Hippocampal response at event boundary* in Methods).

In Figure 4C, the blue line illustrates hippocampal activity at event boundaries with the highest cortical alignment (i.e., score 3) from the *nesting* method. The univariate activity averaged across 2 TRs before an event boundary was subtracted from the BOLD signal at each TR in the graph, following the methods from Lee & Chen (2022) in which the current dataset was originally introduced. A paired *t*-test was performed on hippocampal responses at event boundaries with score 0 (the lowest possible score; 45 boundary trials) and score 3 (the highest possible score; 43 boundary trials) of the nesting method due to the extremely small number of trials in the intervening bins (6 event boundaries for score 1 and 14 event boundaries for score 2). Hippocampal activity during the 5-TR period (0 to 4 TRs) from the onset of an event boundary significantly differed between these two scores, which are at the opposite ends of the score range (*t*(16) = –2.14, *p* = .049; *M* = –0.02, *SD* = 0.07 for score 0; *M* = 0.04, *SD* = 0.06 for score 3).

A one-way repeated measures ANOVA was also performed for each alignment scoring method, this time including the intervening bins (**Supplementary Figure 7**). We found a significant main effect of the cortical alignment score for strict nesting (*F*(4, 64) = 13.86, *p* < .001, 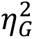 = .39) and nesting (*F*(3, 48) = 10.74, *p* < .001, 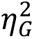 = .32), and a trending effect for summation (*F*(3, 48) = 2.77, *p* = .052, 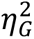 = .12). The hippocampal response was greatest at event boundaries associated with cortical boundary shifts fully aligned across the cortical hierarchy (alignment score = 4 for strict nesting or alignment score = 3 for either nesting or summation). However, while event boundaries with the highest cortical alignment score (blue lines) consistently corresponded to the highest hippocampal response, event boundaries with the lowest cortical alignment score (black lines) did not always elicit the lowest hippocampal response. The results from the post-hoc pairwise comparison are shown in **Supplementary Table 3**. Note that these tests should be treated with caution because of the uneven number of event boundary trials with different alignment scores. Additionally, the observed effects were susceptible to which time window was selected for the baseline: no significant effects were found when using the baseline calculated with a wider time window of 10 TRs, instead of 2 TRs, with the nesting and summation methods. There was a significant main effect with the strict nesting method (*F*(4, 64) = 2.52, *p* = .0495, 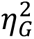 = .12). Overall, while the highest cortical alignment scores did consistently yield the largest hippocampal responses, there did not appear to be strong evidence of a more finely graded hippocampal response based on alignment of pattern shifts across different levels of the cortical hierarchy.

We also examined whether we could observe an event boundary strength modulation effect in the hippocampus, as reported in a prior study (Ben-Yakov & Henson, 2018), using the current dataset. Hippocampal univariate activity was analyzed at event boundaries of different strengths using a one-way repeated measures ANOVA (**Supplementary Figure 8**). We did not replicate prior findings when the baseline activity was calculated using 2 TRs prior to an event boundary, but we observed the numerically highest univariate activity for strong event boundary trials, on average across subjects (*F*(2, 32) = 1.24, *p* > .3, 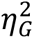 = .06; *M* = 0.02, *SD* = 0.06 for strong; *M* = –0.01, *SD* = 0.04 for moderate; *M* = –0.02, *SD* = 0.08 for weak). When 10-TR baseline was used, a trending main effect of boundary strength was observed (*F*(2, 32) = 3.14, *p* = .057, 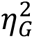 = .14), with the numerically highest hippocampal activity for strong event boundaries as in 2-TR averaging (*M* = 0.04, *SD* = 0.06 for strong; *M* = 0.01, *SD* = 0.06 for moderate; *M* = –0.03, *SD* = 0.10 for weak).

Note that the across-observer agreement range used for defining event boundary strength in the current study was much wider than that of Ben-Yakov and Henson (2018), which presumably affects the results. In the prior study, the event boundaries were identified as being agreed upon by more than five observers, which would fall within the strong category of our event boundaries. Since few or none of the event boundaries in our weak or moderate category were investigated by Ben-Yakov and Henson (2018), a direct comparison is difficult.

## Discussion

In this study, we examined the relationship between event boundary strength and pattern shifts across the cortical hierarchy during naturalistic movie-viewing. Using group-averaged fMRI data, we found that the magnitude of event boundary-triggered multi-voxel pattern shifts (“cortical boundary shifts”) was modulated by event boundary strength, with strong boundaries yielding the largest shifts; this effect was clearer in low-level (short-timescale) sensory areas than in higher-level (longer-timescale) areas. Importantly, for group-level results, apparent gradation could theoretically arise from the fact that strong boundaries have a higher likelihood of detection than moderate or weak boundaries, even while the neural response is “binary”; alternatively, the neural response could be truly “graded”, with larger magnitude responses to stronger boundaries. In order to discriminate between these possibilities, we examined the data at the individual subject level. We observed modulation of cortical pattern shifts by event boundary strength at all levels of the hierarchy, with strong gradation found in auditory cortex. A “binary” response profile should elicit a bimodal distribution of pattern shifts; we did not observe such bimodality at any cortical level. However, it is important to note that while we did not find evidence supporting a binary response profile in any brain region, the current results do not constitute strong evidence *against* the proposal, especially in high-level areas which exhibited weak gradation. Finally, we found that the extent to which cortical boundary shifts occurred simultaneously across different cortical hierarchical levels was associated with event boundary strength and hippocampal responses at event boundaries.

### Granularity of event segmentation and event boundary strength

Event Segmentation Theory (EST; Zacks et al., 2007) proposes that the representations in event models are multi-modal, and our continuous experience can be segmented at multiple timescales simultaneously. When two groups of individuals perform event segmentation at different levels of granularity, coarse-grained and fine-grained event boundaries are temporally aligned (Zacks, Tversky, et al., 2001); this has been suggested as evidence supporting the idea of multi-timescale event segmentation. In behavioral studies, the granularity of event segmentation was suggested to reflect processing timescales. On the other hand, recent research has explored hippocampal responses at event boundaries with varying salience or strengths by allowing participants to segment movies at the granularity that they feel is natural, and by aggregating these event boundaries from different individuals (Ben-Yakov & Henson, 2018). However, it is challenging to disentangle the concepts of granularity in event segmentation and event boundary strength. In the current study, we sought to verify whether agreement across individuals corresponds to event boundary strength; after obtaining timestamps of event boundaries with different degrees of boundary agreement across observers following the methods of Ben-Yakov and Henson (2018), we asked an independent group of people to rate the strength of those pre-identified event boundaries. We observed a strong positive relationship between event boundary agreement and strength ratings. Future studies might investigate how segmentation granularity relates to both agreement and perceived strength of boundaries, and to what extent fine-grained event boundaries defined from human behavior relate to cortical pattern shifts in short-timescale sensory areas.

### Changes in perceptual and situational features and event segmentation

Prior studies have revealed that event boundaries are somewhat normative: the behavior of event segmentation is fairly consistent and reliable between young and older adults (Kurby & Zacks, 2018; for coarse segmentation) as well as across people in a relatively homogeneous age group (Hanson & Hirst, 1989), despite the inherent individual differences in event segmentation that are stable over more than a year (Speer et al., 2003). According to EST (Zacks et al., 2007), people perceive an event boundary when the prediction of the immediate future is violated by the actual outcome. Compatible with this account, it has been shown that even reward prediction errors can create a new latent state, thereby disrupting integration of information across distinct reward states (Rouhani et al., 2020). In narratives, it has been suggested that one or more changes in perceptual or situational features in ongoing activity are key to leading to such prediction errors. Empirical work has shown that the moments of perceptual or situational changes, such as locations (spatial context), time (temporal context), goals, characters, objects and movements, were associated with event boundaries (Speer et al., 2007; Zacks, Kumar, et al., 2009). Also, the number of changes in these features significantly predicted the probability of being indicated as an event boundary for narrative texts or films (Zacks et al., 2010; Zacks, Speer, et al., 2009). Neuroimaging studies further revealed neural correlates of event boundaries, which involved shifts in perceptual or situational features by experimental manipulations (Whitney et al., 2009; Zacks et al., 2010).

In the current study, an audiovisual narrative was used as a stimulus, and subjects passively watched a series of movies in the scanner. In these settings, it is likely that *spontaneous* event segmentation relies heavily on detecting perceptual changes such as film cuts, which entail shifts in situational features such as locations, time, and characters (note that fMRI subjects did not perform an event segmentation task). In both group-level and individual-level analyses, we consistently observed that a graded response profile is pronounced in low-level sensory processing areas with short timescales, whereas magnitude differences in cortical boundary shifts between different event boundary strengths are less discernible in high-level (longer-timescale) areas. We speculate that, in our data, weak event boundaries might be associated with perceptual changes such as slight changes in motion, facial expressions, or camera angle, while strong event boundaries might occur at moments of both perceptual and situational changes in multiple dimensions, as behavioral studies have revealed the relationship between the number of perceptual and situational changes and the probability of event segmentation (Zacks et al., 2010; Zacks, Speer, et al., 2009).

### Brain response to event boundaries: univariate vs. multivariate analysis

In the current study, we investigated cortical response profiles at the moments when people perceive event boundaries in an ongoing experience. Based on findings from recent research on neural event segmentation processes (Baldassano et al., 2017, Geerligs et al., 2021), we tracked changes in multi-voxel patterns elicited by naturalistic movie-viewing. Earlier neuroimaging studies have investigated univariate responses to coarse-grained vs. fine-grained boundaries in movies and text narratives, with somewhat mixed results. In the left and right posterior inferior temporal sulcus and the left fusiform gyrus, greater responses were found at coarse-grained boundaries than at fine-grained boundaries while participants passively watched short video clips of daily activities, such as making a bed (Zacks, Braver, et al., 2001). In another study, univariate activity in the precuneus, a core region within the DMN, appeared to be more positive at coarse-grained event boundaries than at fine-grained ones when participants read written stories in the fMRI scanner (Speer et al., 2007). In contrast, during cinematic movie-viewing, a different pattern was observed in bilateral regions at the parietal-temporal-occipital junction, which encompasses the angular gyrus, another core DMN region (Zacks et al., 2010); univariate activity was greater at fine-grained event boundaries compared to coarse-grained ones. Is the multi-voxel pattern-based neural event segmentation phenomenon essentially equivalent to the univariate brain response to event boundaries discovered by earlier studies? If not, do they reflect different aspects of event segmentation? Further research is needed to elucidate the relationship between univariate and pattern shift responses at event boundaries.

### Information integrated at different timescales and neural event segmentation

In prior studies which examined multi-voxel pattern based neural event segmentation, a cortical region’s information processing timescale was measured by the overall duration of neural states in that region evoked during movie-viewing (Baldassano et al., 2017; Geerligs et al., 2022). In our study, we opted to define the cortical hierarchy of timescales by calculating temporal receptive windows (TRWs). This methodological choice is grounded in numerous prior studies demonstrating that a cortical area’s processing timescale can robustly be assessed by measuring the extent to which the region’s response to current input is influenced by past information (Hasson et al., 2008; Lerner et al., 2011). For example, if a brain region’s response is undisrupted by changes in the external stimulus a second ago, that area is deemed to have a timescale of less than a second (e.g., early sensory processing areas). In a cortical region with a long timescale, responses at the current moment are sensitive to changes that occurred tens of seconds ago; high-level cortical areas with long timescales can retain information for around 30 seconds during natural continuous stimulus processing (Hasson et al., 2015), even in the absence of the hippocampus (Zuo et al., 2020). In the DMN, which has long timescales, we observed a weak but significant modulation of multi-voxel pattern changes by boundary strength: multi-voxel patterns from immediately before and after an event boundary were more distinct at strong boundaries compared to weak or moderate ones. This might suggest that the DMN can rapidly respond to event boundaries in a strength-sensitive manner despite its long processing timescale, consistent with prior findings that high-level, long-timescale cortical areas can forget prior context at a rate similar to that in low-level brain areas (Chien & Honey, 2020).

### Information flow in the cortex during event segmentation

Intriguingly, recent research has examined how stimulus information is transmitted from lower to higher levels along the cortical hierarchy of timescales: functional coupling between early auditory processing areas and DMN regions exhibits a temporal lag during audio narrative listening (Chang et al., 2022). This effect disappeared when participants listened to a temporally scrambled, hardly interpretable story. One observation in our study was that most event boundaries in the “strong” category were paired with the highest cortical alignment score, regardless of the scoring method. This raises the question of how variation in perceived boundary strength is linked to information transmission from low-level to high-level cortical areas (e.g., detecting changes in perceptual information). Event boundaries can be driven by sensory changes in a stimulus (i.e., “bottom-up”) as well as by more abstract transitions in ongoing experience. For example, people perceive a boundary when the task goal switches (Y. C. Wang & Egner, 2022), and when they encounter a transition point in stimulus based on a learned temporal structure in stimulus presentation (Schapiro et al., 2013). In our study, we speculate that the event boundaries were mostly driven by sensory changes in the movie, and under these conditions the bottom-up information flow from sensory regions to high-level cortical areas within the DMN may be the dominant process. An interesting possibility is that, in experiences that are more self-initiated such as spoken recollection or conversation, “top-down” event boundaries would have an inverted alignment structure: alignment would be measured starting from the highest hierarchical level and “nested” downward, and boundary strength would scale with the degree of this alignment.

In summary, in cortical areas at multiple timescales, we observed that multi-voxel pattern shifts at event boundaries were modulated by boundary strength during naturalistic movie-viewing. There appeared to be differentiation across the cortical hierarchy: auditory processing areas with short timescales exhibited a strongly graded response, whereas modulation by boundary strength was present, but weaker, in areas with longer timescales such as in the DMN. Alignment of boundary-related pattern shifts across multiple levels of the cortical hierarchy was correlated with perceived boundary strength as well. These observations add to our understanding of how event boundaries are instantiated in brain activity during naturalistic experiences.

## Acknowledgments

We thank Dr. Christopher Honey for valuable intellectual input on designing analyses and for discussions; Savannah Born for help with data collection.

## Data Availability Statement

Data will be made available upon publication.

## Author Contributions

Yoonjung Lee: Conceptualization; Data Curation; Formal Analysis; Investigation; Methodology; Project Administration; Software; Visualization; Writing – Original Draft Preparation; Writing – Review & Editing. Janice Chen: Conceptualization; Funding acquisition; Investigation; Methodology; Resources; Software; Supervision; Writing – Original Draft Preparation; Writing – Review & Editing

## Diversity in Citation Practices

Recent work in several fields of science has identified a bias in citation practices such that papers from women and other minority scholars are under-cited relative to the number of such papers in the field (Bertolero et al., 2020; Caplar et al., 2017; Chatterjee & Werner, 2021; Dion et al., 2018; Dworkin et al., 2020; Fulvio et al., 2021; Maliniak et al., 2013; Mitchell et al., 2013; X. Wang et al., 2021). Here we sought to proactively consider choosing references that reflect the diversity of the field in thought, form of contribution, gender, race, ethnicity, and other factors. First, we obtained the predicted gender of the first and last author of each reference by using databases that store the probability of a first name being carried by a woman (Dworkin et al., 2020; Zhou et al., 2020). By this measure and excluding self-citations to the first and last authors of our current paper), our references contain 5.41% woman(first)/woman(last), 18.92% man/woman, 37.84% woman/man, and 37.84% man/man. This method is limited in that (a) names, pronouns, and social media profiles used to construct the databases may not, in every case, be indicative of gender identity and (b) it cannot account for intersex, non-binary, or transgender people. Second, we obtained predicted racial/ethnic category of the first and last author of each reference by databases that store the probability of a first and last name being carried by an author of color (Ambekar et al., 2009; Chintalapati et al., 2018). By this measure (and excluding self-citations), our references contain 2.92% author of color (first)/author of color(last), 19.38% white author/author of color, 17.48% author of color/white author, and 60.22% white author/white author. This method is limited in that (a) names and Florida Voter Data to make the predictions may not be indicative of racial/ethnic identity, and (b) it cannot account for Indigenous and mixed-race authors, or those who may face differential biases due to the ambiguous racialization or ethnicization of their names. We look forward to future work that could help us to better understand how to support equitable practices in science.

## SUPPLEMENTARY INFORMATION

**Supplementary Table 1.** Agreement range for each event boundary strength category.

|  | Proportion of human observers |  |
| --- | --- | --- |
|  | Minimum | Maximum |
| Weak | .11 | .19 |
| Moderate | .20 | .40 |
| Strong | .40 | .98 |

**Supplementary Table 2.** Agreement range for each event boundary strength category after excluding pairs of event boundaries less than 6 seconds apart.

|  | Proportion of human observers |  |
| --- | --- | --- |
|  | Minimum | Maximum |
| Weak | .12 | .19 |
| Moderate | .20 | .38 |
| Strong | .40 | .98 |

**Supplementary Figure 1.**
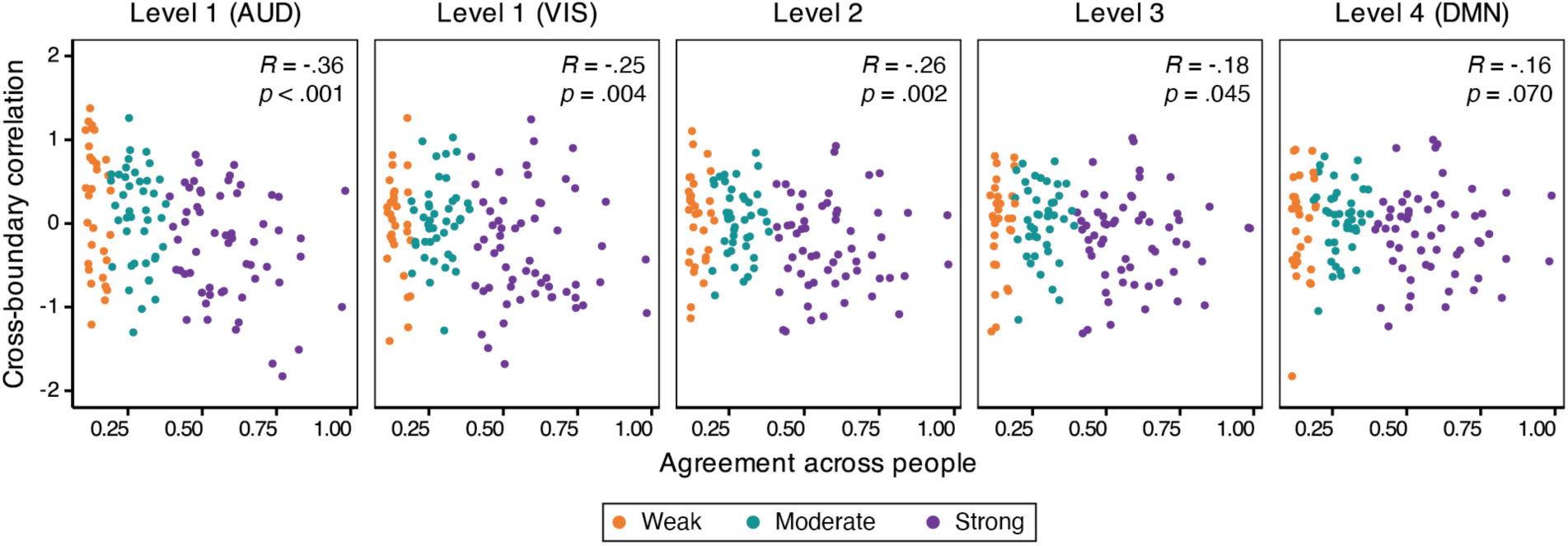
Spearman’s rank correlation between cortical pattern shift and across-observer boundary agreement at each cortical hierarchical level. Note that correlation time series was normalized (see Methods).

**Supplementary Figure 2.**
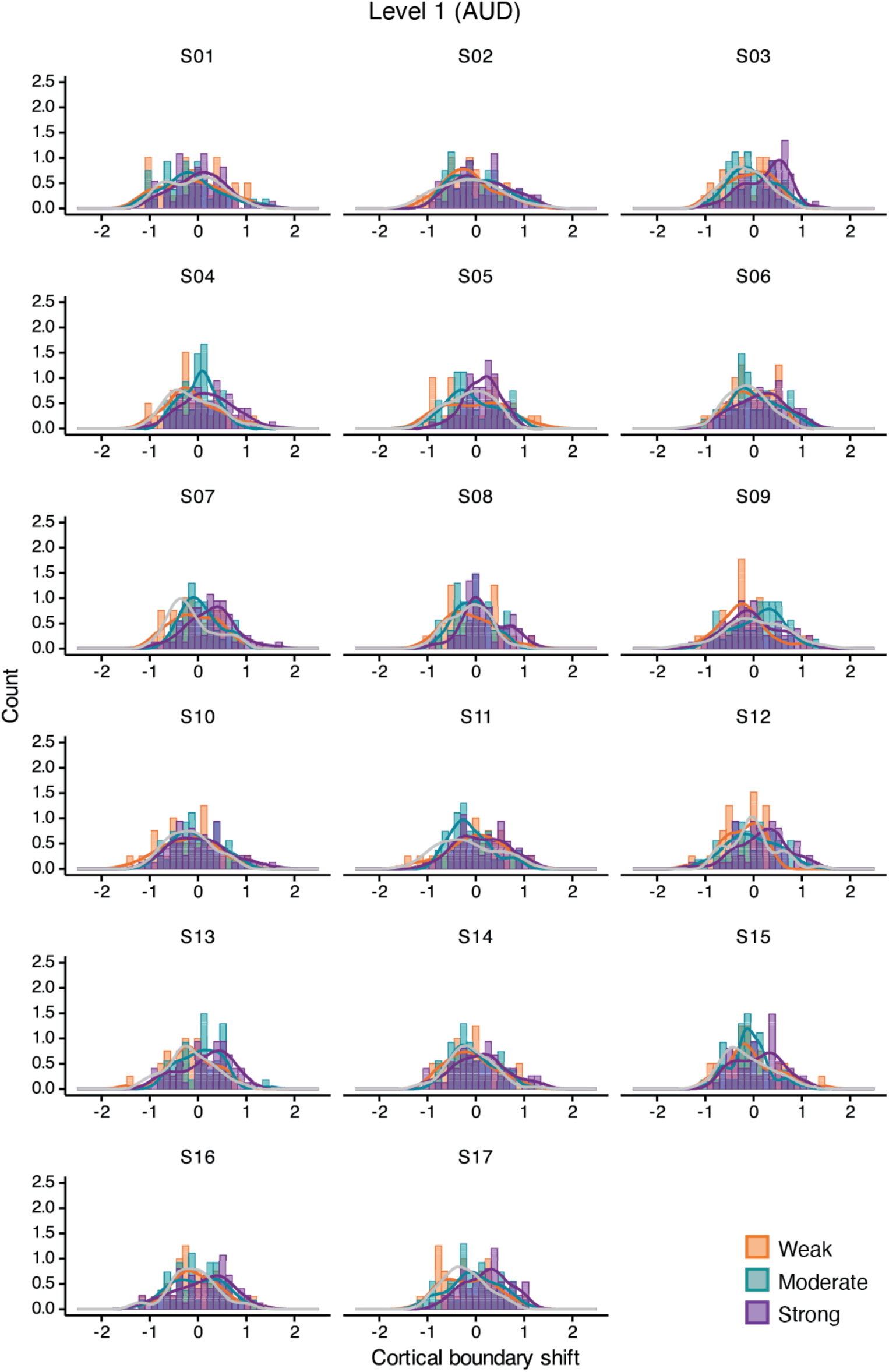
Individuals’ distributions of cortical boundary shifts for each boundary strength category in auditory processing areas (AUD) at level 1 of the cortical hierarchy. The gray line depicts a density curve of the non-boundary data.

**Supplementary Figure 3.**
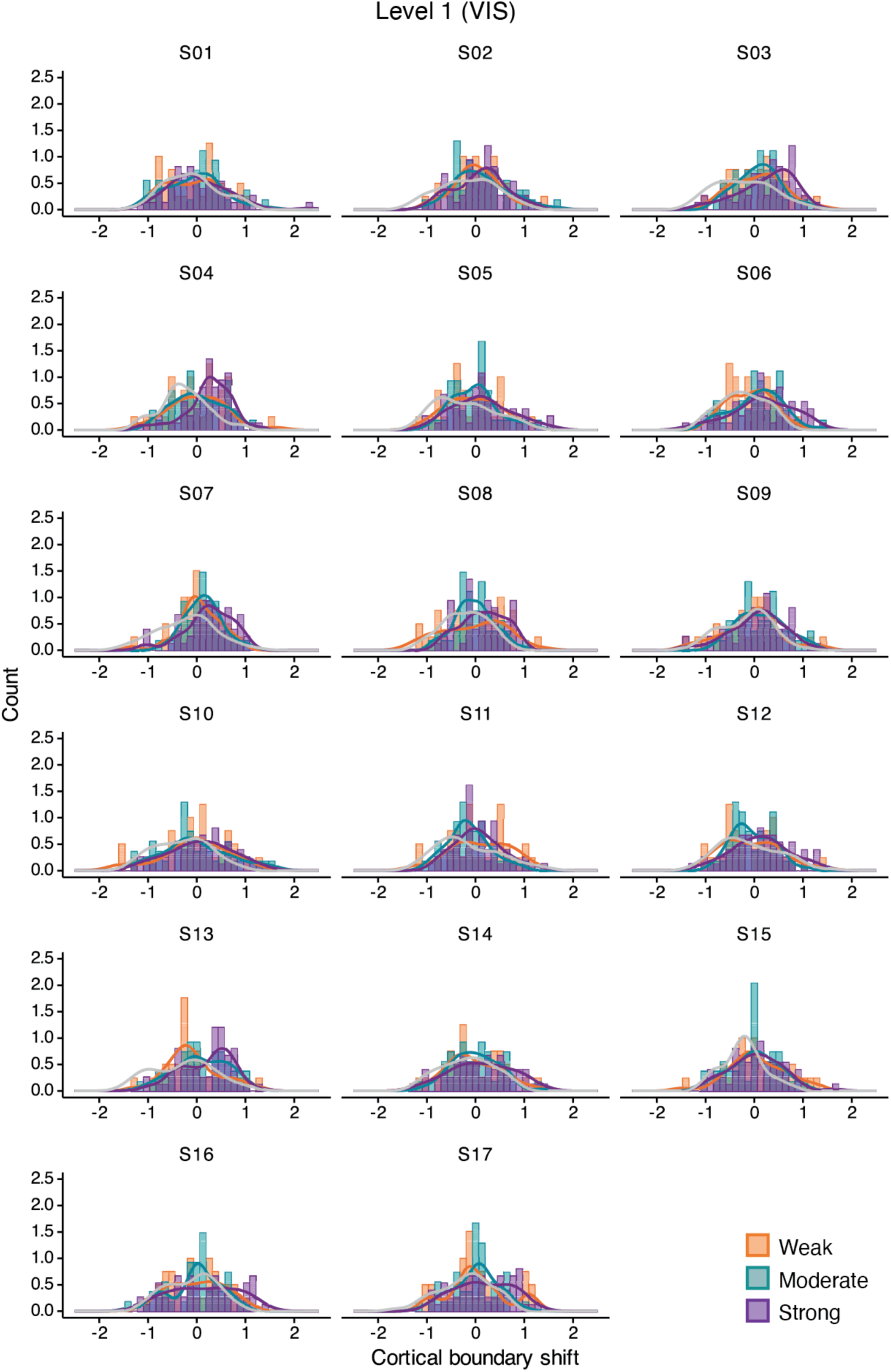
Individuals’ distributions of cortical boundary shifts for each boundary strength category in visual processing areas (VIS) at level 1 of the cortical hierarchy. The gray line depicts a density curve of the non-boundary data.

**Supplementary Figure 4.**
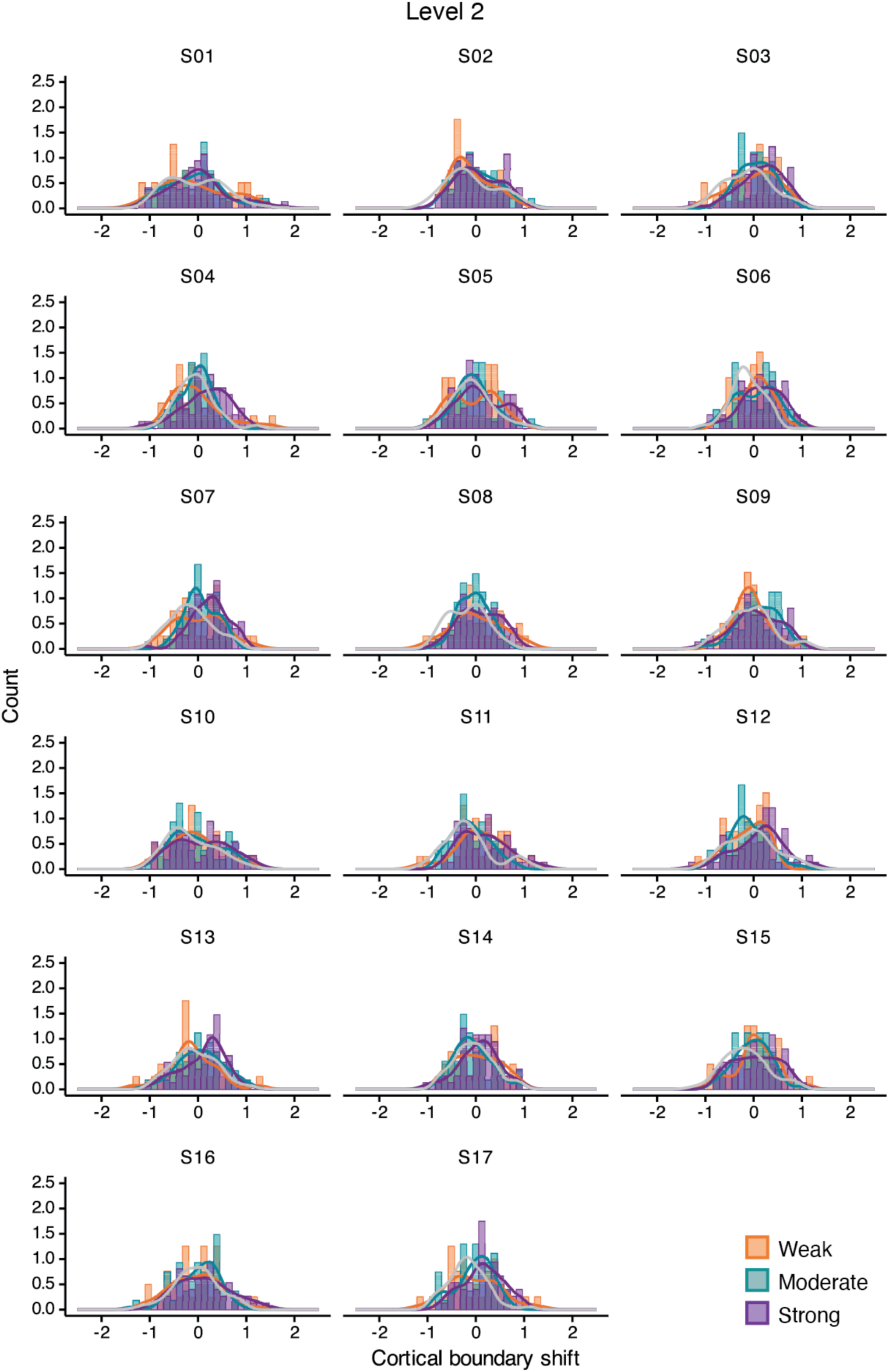
Individuals’ distributions of cortical boundary shifts for each boundary strength category in the areas at level 2 of the cortical hierarchy. The gray line depicts a density curve of the non-boundary data.

**Supplementary Figure 5.**
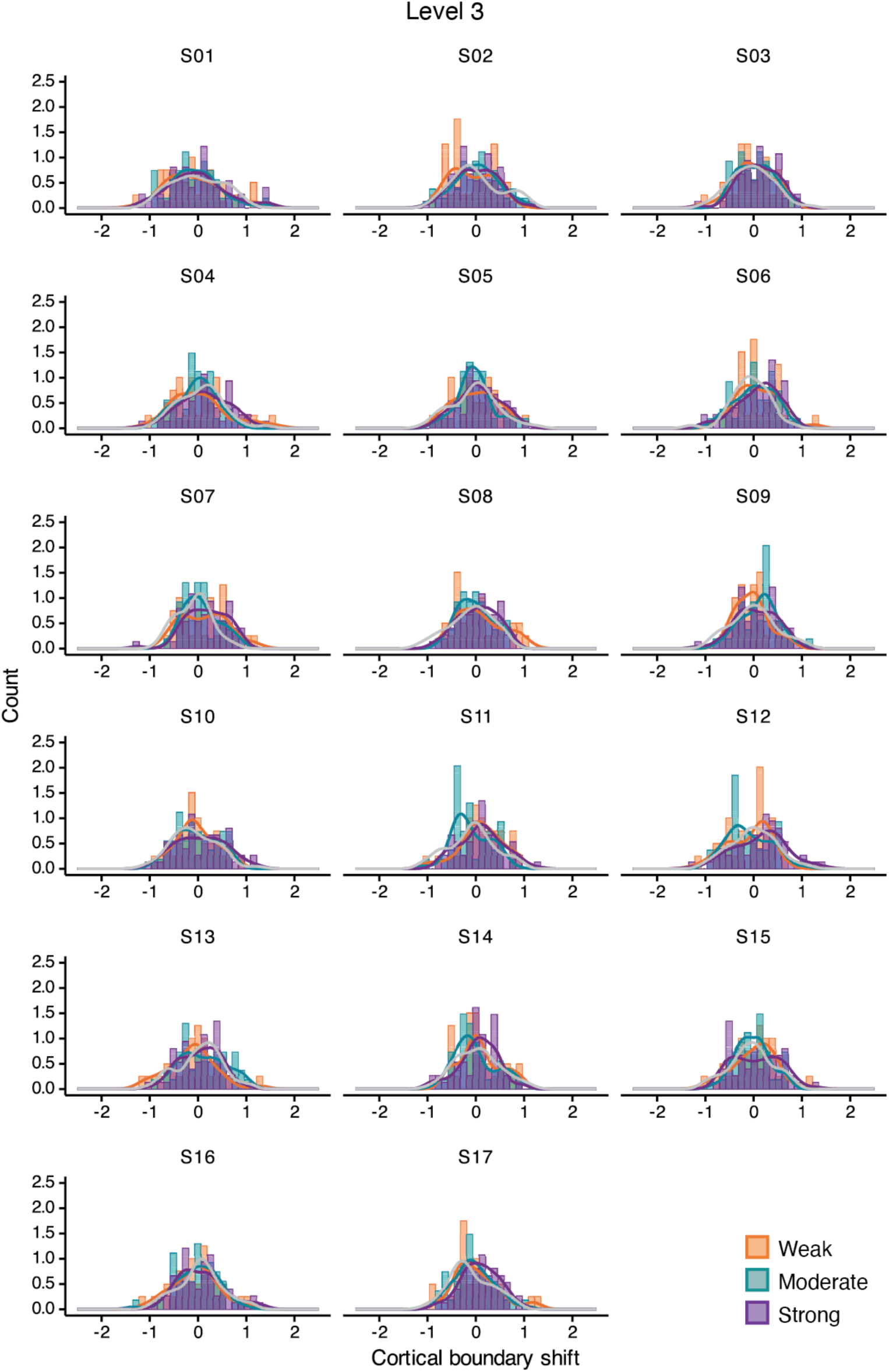
Individuals’ distributions of cortical boundary shifts for each boundary strength category in the areas at level 3 of the cortical hierarchy. The gray line depicts a density curve of the non-boundary data.

**Supplementary Figure 6.**
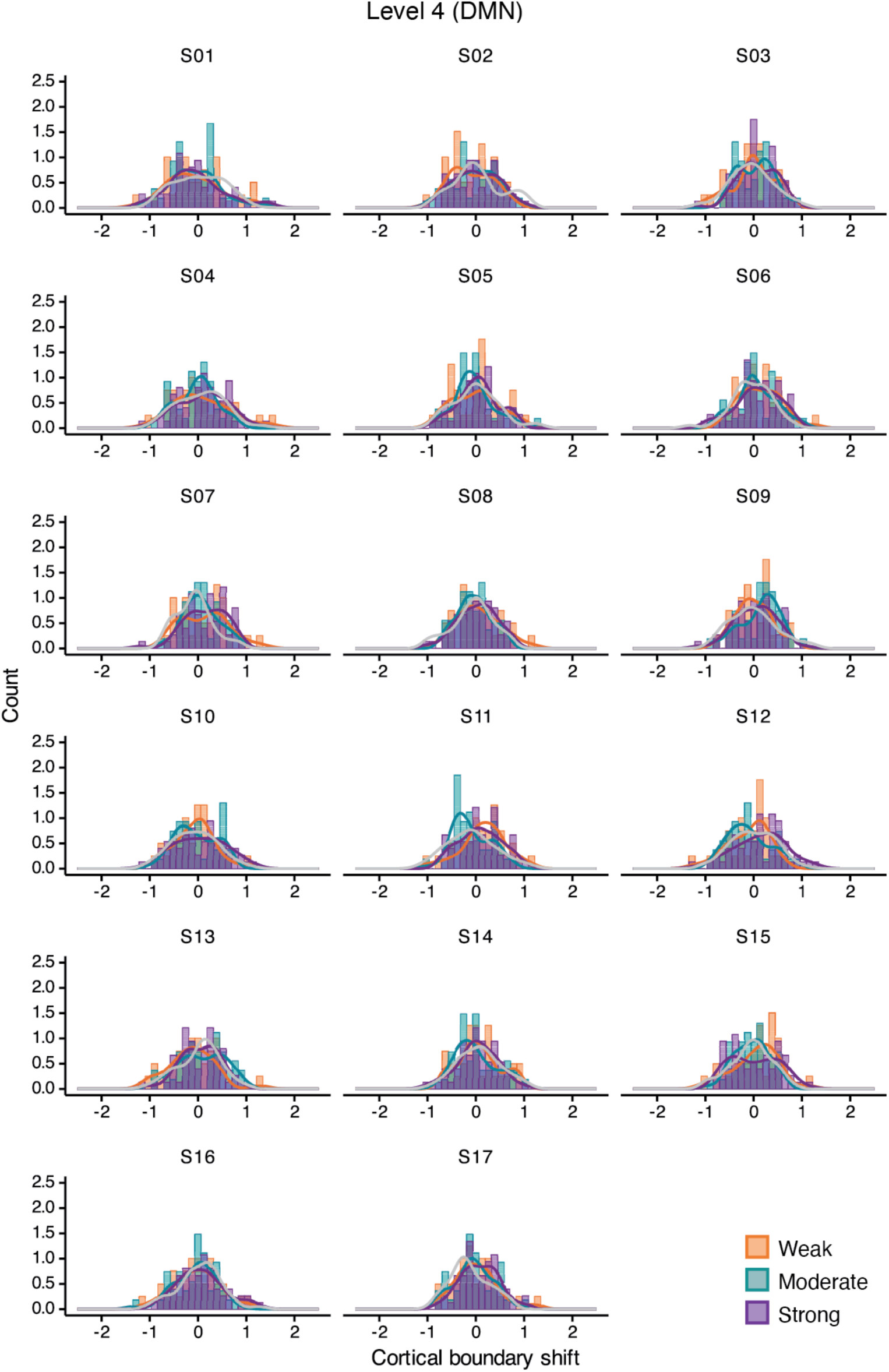
Individuals’ distributions of cortical boundary shifts for each boundary strength category in the default mode network (DMN) areas at level 4 of the cortical hierarchy. The gray line depicts a density curve of the non-boundary data.

**Supplementary Figure 7.**
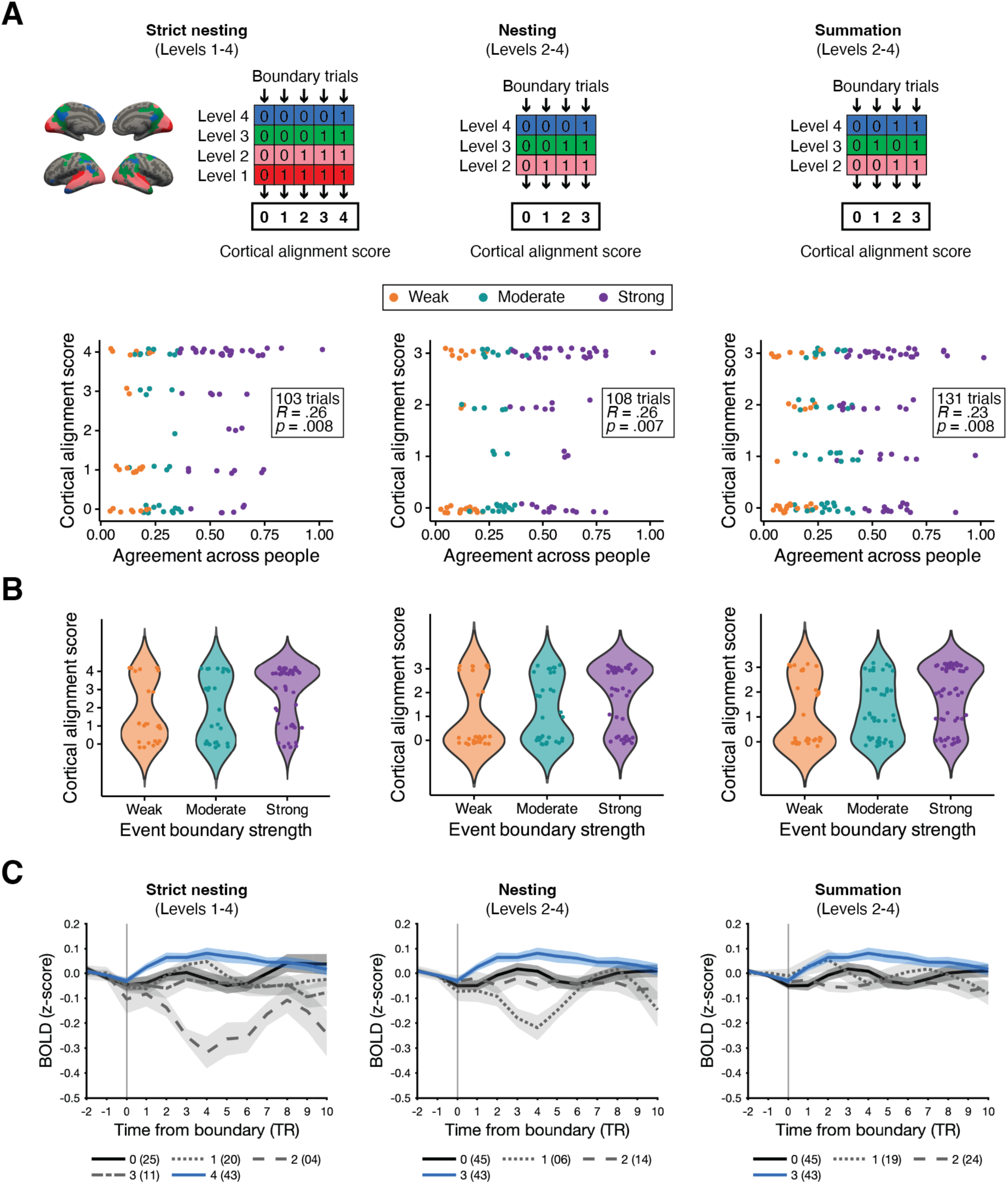
Three types of scoring methods (strict nesting, nesting, and summation). Due to the absence of a bottom-up nested structure in cortical pattern shifts, 21.4% and 17.6% of event boundary trials were excluded for strict nesting and nesting, respectively. (A) There was a significant positive relationship between the degree of cortical alignment and boundary agreement across people for all scoring methods (Spearman’s rank correlation). In the scatter plot, individual dots depict event boundary trials. (B) The same data are presented in a violin plot to visualize the distribution. (C) Each line illustrates hippocampal activity at event boundaries with different alignment scores for each scoring method (blue solid line: the highest possible alignment score; black solid line: the lowest possible alignment score; shaded area: ± SEM across subjects). Different line types depict different scores (gray dotted line: score 1, gray dashed line: score 2, gray dash-dotted line in strict nesting: score 3). In the legend, the number in a parenthesis shows the number of event boundary trials associated with that score. Note that different scoring methods produce different numbers of boundary trials for a given score.

**Supplementary Table 3.**
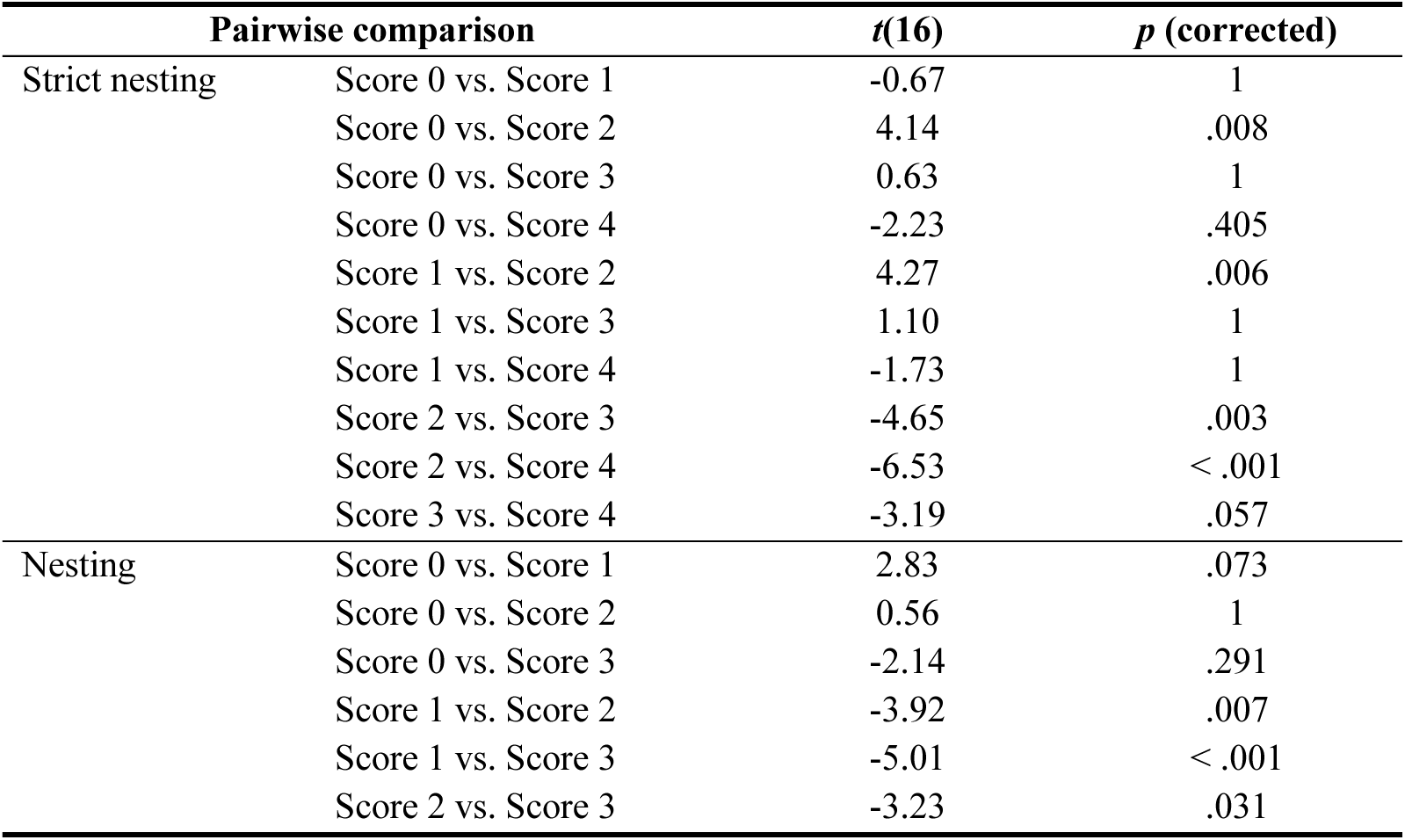
Post-hoc pairwise comparison (paired *t*-test) results in the hippocampus analysis. Bonferroni correction was applied. The corrected *p*-value is the initial *p*-value multiplied by the number of comparisons. Note that no significant main effect of cortical alignment was observed with the summation method.

**Supplementary Figure 8.**
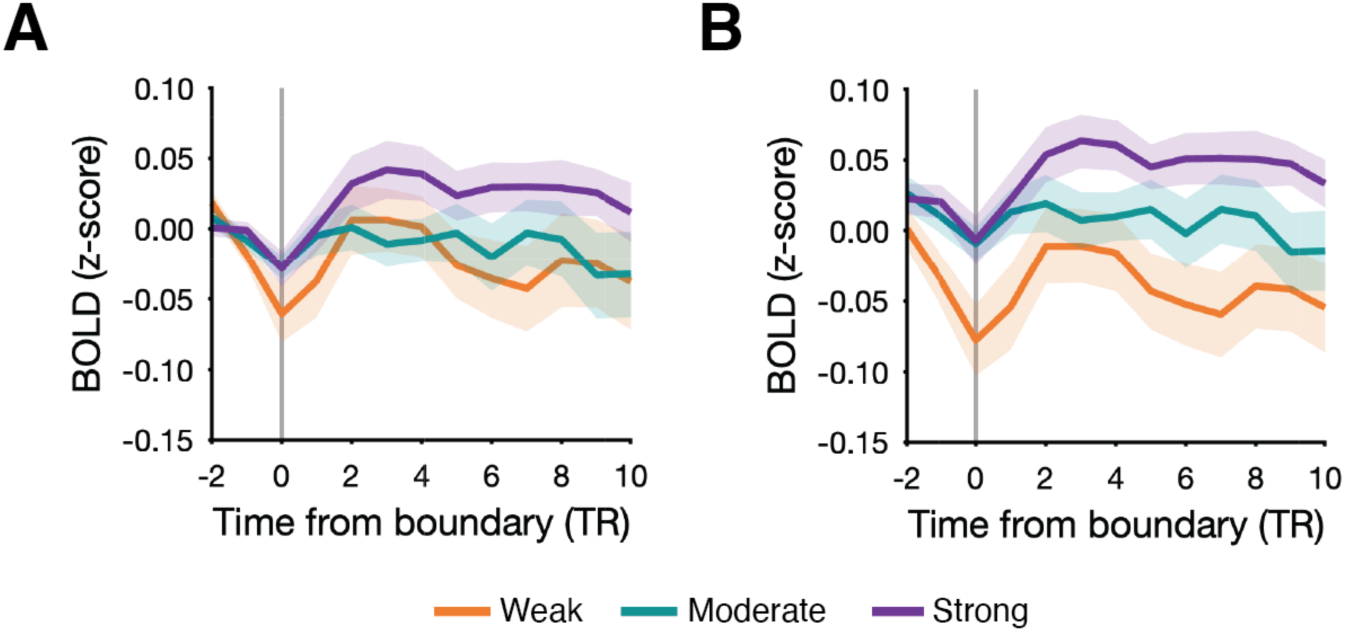
Hippocampal response at event boundaries varying with strengths.

## References

1. Ambekar, A., Ward, C., Mohammed, J., Male, S., & Skiena, S. (2009). Name-ethnicity classification from open sources. Proceedings of the 15th ACM SIGKDD International Conference on Knowledge Discovery and Data Mining, 49–58. 10.1145/1557019.1557032

2. Baldassano, C., Chen, J., Zadbood, A., Pillow, J. W., Hasson, U., & Norman, K. A. (2017). Discovering Event Structure in Continuous Narrative Perception and Memory. Neuron, 95(3), 709–721.e5. 10.1016/j.neuron.2017.06.041

3. Ben-Yakov, A., & Dudai, Y. (2011). Constructing Realistic Engrams: Poststimulus Activity of Hippocampus and Dorsal Striatum Predicts Subsequent Episodic Memory. The Journal of Neuroscience, 31(24), 9032. 10.1523/JNEUROSCI.0702-11.2011

4. Ben-Yakov, A., Eshel, N., & Dudai, Y. (2013). Hippocampal immediate poststimulus activity in the encoding of consecutive naturalistic episodes. Journal of Experimental Psychology: General, 142(4), 1255–1263. 10.1037/a0033558

5. Ben-Yakov, A., & Henson, R. N. (2018). The Hippocampal Film Editor: Sensitivity and Specificity to Event Boundaries in Continuous Experience. Journal of Neuroscience, 38(47), 10057–10068. 10.1523/JNEUROSCI.0524-18.2018

6. Bertolero, M. A., Dworkin, J. D., David, S. U., Lloreda, C. L., Srivastava, P., Stiso, J., Zhou, D., Dzirasa, K., Fair, D. A., Kaczkurkin, A. N., Marlin, B. J., Shohamy, D., Uddin, L. Q., Zurn, P., & Bassett, D. S. (2020). Racial and ethnic imbalance in neuroscience reference lists and intersections with gender. BioRxiv, 2020–10. 10.1101/2020.10.12.336230

7. Caplar, N., Tacchella, S., & Birrer, S. (2017). Quantitative evaluation of gender bias in astronomical publications from citation counts. Nature Astronomy, 1(6), 0141. 10.1038/s41550-017-0141

8. Chang, C. H. C., Nastase, S. A., & Hasson, U. (2022). Information flow across the cortical timescale hierarchy during narrative construction. Proceedings of the National Academy of Sciences, 119(51), e2209307119. 10.1073/pnas.2209307119

9. Chatterjee, P., & Werner, R. M. (2021). Gender Disparity in Citations in High-Impact Journal Articles. JAMA Network Open, 4(7), e2114509–e2114509. 10.1001/jamanetworkopen.2021.14509

10. Chen, J., Honey, C. J., Simony, E., Arcaro, M. J., Norman, K. A., & Hasson, U. (2016). Accessing Real-Life Episodic Information from Minutes versus Hours Earlier Modulates Hippocampal and High-Order Cortical Dynamics. Cerebral Cortex, 26(8), 3428–3441. 10.1093/cercor/bhv155

11. Chien, H.-Y. S., & Honey, C. J. (2020). Constructing and Forgetting Temporal Context in the Human Cerebral Cortex. Neuron, 106(4), 675–686.e11. 10.1016/j.neuron.2020.02.013

12. Chintalapati, R., Laohaprapanon, S., & Sood, G. (2018). Predicting Race and Ethnicity From the Sequence of Characters in a Name. arXiv preprint arXiv:1805.02109. 10.48550/arXiv.1805.02109

13. Desikan, R. S., Ségonne, F., Fischl, B., Quinn, B. T., Dickerson, B. C., Blacker, D., Buckner, R. L., Dale, A. M., Maguire, R. P., Hyman, B. T., Albert, M. S., & Killiany, R. J. (2006). An automated labeling system for subdividing the human cerebral cortex on MRI scans into gyral based regions of interest. NeuroImage, 31(3), 968–980. 10.1016/j.neuroimage.2006.01.021

14. Dion, M. L., Mitchell, S. M., & Sumner, J. L. (2018). Gendered Citation Patterns across Political Science and Social Science Methodology Fields. Political Analysis, 26(3), 312–327. 10.1017/pan.2018.12

15. DuBrow, S., & Davachi, L. (2013). The influence of context boundaries on memory for the sequential order of events. Journal of Experimental Psychology: General, 142(4), 1277–1286. APA PsycArticles. 10.1037/a0034024

16. DuBrow, S., & Davachi, L. (2016). Temporal binding within and across events. Neurobiology of Learning and Memory, 134, 107–114. 10.1016/j.nlm.2016.07.011

17. Dworkin, J. D., Linn, K. A., Teich, E. G., Zurn, P., Shinohara, R. T., & Bassett, D. S. (2020). The extent and drivers of gender imbalance in neuroscience reference lists. Nature Neuroscience, 23(8), 918–926. 10.1038/s41593-020-0658-y

18. Ezzyat, Y., & Davachi, L. (2011). What Constitutes an Episode in Episodic Memory? Psychological Science, 22(2), 243–252. 10.1177/0956797610393742

19. Ezzyat, Y., & Davachi, L. (2014). Similarity Breeds Proximity: Pattern Similarity within and across Contexts Is Related to Later Mnemonic Judgments of Temporal Proximity. Neuron, 81(5), 1179– 1189. 10.1016/j.neuron.2014.01.042

20. Fulvio, J. M., Akinnola, I., & Postle, B. R. (2021). Gender (Im)balance in Citation Practices in Cognitive Neuroscience. Journal of Cognitive Neuroscience, 33(1), 3–7. 10.1162/jocn_a_01643

21. Geerligs, L., Gözükara, D., Oetringer, D., Campbell, K. L., van Gerven, M., & Güçlü, U. (2022). A partially nested cortical hierarchy of neural states underlies event segmentation in the human brain. eLife, 11, e77430. 10.7554/eLife.77430

22. Geerligs, L., van Gerven, M., & Güçlü, U. (2021). Detecting neural state transitions underlying event segmentation. NeuroImage, 236, 118085. 10.1016/j.neuroimage.2021.118085

23. Hanson, C., & Hirst, W. (1989). On the representation of events: A study of orientation, recall, and recognition. Journal of Experimental Psychology: General, 118(2), 136–147. 10.1037/0096-3445.118.2.136

24. Hartigan, J. A., & Hartigan, P. M. (1985). The Dip Test of Unimodality. The Annals of Statistics, 13(1), 70–84. JSTOR. 10.1214/aos/1176346577

25. Hasson, U., Chen, J., & Honey, C. J. (2015). Hierarchical process memory: Memory as an integral component of information processing. Trends in Cognitive Sciences, 19(6), 304–313. 10.1016/j.tics.2015.04.006

26. Hasson, U., Yang, E., Vallines, I., Heeger, D. J., & Rubin, N. (2008). A Hierarchy of Temporal Receptive Windows in Human Cortex. The Journal of Neuroscience, 28(10), 2539. 10.1523/JNEUROSCI.5487-07.2008

27. Heusser, A. C., Ezzyat, Y., Shiff, I., & Davachi, L. (2018). Perceptual boundaries cause mnemonic trade-offs between local boundary processing and across-trial associative binding. *Journal of Experimental Psychology: Learning*, Memory, and Cognition, 44, 1075–1090. 10.1037/xlm0000503

28. JASP Team (2024). JASP (Version 0.18.3) [Computer software]

29. Kurby, C. A., & Zacks, J. M. (2018). Preserved neural event segmentation in healthy older adults. Psychology and Aging, 33(2), 232–245. 10.1037/pag0000226

30. Lee, H., & Chen, J. (2022). Predicting memory from the network structure of naturalistic events. Nature Communications, 13(1), 4235. 10.1038/s41467-022-31965-2

31. Lerner, Y., Honey, C. J., Silbert, L. J., & Hasson, U. (2011). Topographic Mapping of a Hierarchy of Temporal Receptive Windows Using a Narrated Story. The Journal of Neuroscience, 31(8), 2906. 10.1523/JNEUROSCI.3684-10.2011

32. Maliniak, D., Powers, R., & Walter, B. F. (2013). The Gender Citation Gap in International Relations. International Organization, 67(4), 889–922. 10.1017/S0020818313000209

33. Michelmann, S., Hasson, U., & Norman, K. A. (2023). Evidence That Event Boundaries Are Access Points for Memory Retrieval. Psychological Science, 34(3), 326–344. 10.1177/09567976221128206

34. Michelmann, S., Price, A. R., Aubrey, B., Strauss, C. K., Doyle, W. K., Friedman, D., Dugan, P. C., Devinsky, O., Devore, S., Flinker, A., Hasson, U., & Norman, K. A. (2021). Moment-by-moment tracking of naturalistic learning and its underlying hippocampo-cortical interactions. Nature Communications, 12(1), 5394. 10.1038/s41467-021-25376-y

35. Mitchell, S. M., Lange, S., & Brus, H. (2013). Gendered Citation Patterns in International Relations Journals1. International Studies Perspectives, 14(4), 485–492. 10.1111/insp.12026

36. Newtson, D. (1973). Attribution and the unit of perception of ongoing behavior. Journal of Personality and Social Psychology, 28(1), 28–38. 10.1037/h0035584

37. Rouhani, N., Norman, K. A., Niv, Y., & Bornstein, A. M. (2020). Reward prediction errors create event boundaries in memory. Cognition, 203, 104269. 10.1016/j.cognition.2020.104269

38. Schaefer, A., Kong, R., Gordon, E. M., Laumann, T. O., Zuo, X.-N., Holmes, A. J., Eickhoff, S. B., & Yeo, B. T. T. (2018). Local-Global Parcellation of the Human Cerebral Cortex from Intrinsic Functional Connectivity MRI. Cerebral Cortex, 28(9), 3095–3114. 10.1093/cercor/bhx179

39. Schapiro, A. C., Rogers, T. T., Cordova, N. I., Turk-Browne, N. B., & Botvinick, M. M. (2013). Neural representations of events arise from temporal community structure. Nature Neuroscience, 16(4), 486–492. 10.1038/nn.3331

40. Speer, N. K., Swallow, K. M., & Zacks, J. M. (2003). Activation of human motion processing areas during event perception. *Cognitive, Affective*, & Behavioral Neuroscience, 3(4), 335–345. 10.3758/CABN.3.4.335

41. Speer, N. K., Zacks, J. M., & Reynolds, J. R. (2007). Human Brain Activity Time-Locked to Narrative Event Boundaries. Psychological Science, 18(5), 449–455. 10.1111/j.1467-9280.2007.01920.x

42. Wang, L., Mruczek, R. E. B., Arcaro, M. J., & Kastner, S. (2015). Probabilistic Maps of Visual Topography in Human Cortex. Cerebral Cortex, 25(10), 3911–3931. 10.1093/cercor/bhu277

43. Wang, X., Dworkin, J. D., Zhou, D., Stiso, J., Falk, E. B., Bassett, D. S., Zurn, P., & Lydon-Staley, D. M. (2021). Gendered citation practices in the field of communication. Annals of the International Communication Association, 45(2), 134–153. 10.1080/23808985.2021.1960180

44. Wang, Y. C., & Egner, T. (2022). Switching task sets creates event boundaries in memory. Cognition, 221, 104992. 10.1016/j.cognition.2021.104992

45. Whitney, C., Huber, W., Klann, J., Weis, S., Krach, S., & Kircher, T. (2009). Neural correlates of narrative shifts during auditory story comprehension. NeuroImage, 47(1), 360–366. 10.1016/j.neuroimage.2009.04.037

46. Zacks, J. M., Braver, T. S., Sheridan, M. A., Donaldson, D. I., Snyder, A. Z., Ollinger, J. M., Buckner, R. L., & Raichle, M. E. (2001). Human brain activity time-locked to perceptual event boundaries. Nature Neuroscience, 4(6), 651–655. 10.1038/88486

47. Zacks, J. M., Kumar, S., Abrams, R. A., & Mehta, R. (2009). Using movement and intentions to understand human activity. Cognition, 112(2), 201–216. 10.1016/j.cognition.2009.03.007

48. Zacks, J. M., Speer, N. K., & Reynolds, J. R. (2009). Segmentation in reading and film comprehension. Journal of Experimental Psychology: General, 138(2), 307–327. 10.1037/a0015305

49. Zacks, J. M., Speer, N. K., Swallow, K. M., Braver, T. S., & Reynolds, J. R. (2007). Event perception: A mind-brain perspective. Psychological Bulletin, 133(2), 273–293. 10.1037/0033-2909.133.2.273

50. Zacks, J. M., Speer, N., Swallow, K., & Maley, C. (2010). The Brain’s Cutting-Room Floor: Segmentation of Narrative Cinema. Frontiers in Human Neuroscience, 4, 168. 10.3389/fnhum.2010.00168

51. Zacks, J. M., Tversky, B., & Iyer, G. (2001). Perceiving, remembering, and communicating structure in events. Journal of Experimental Psychology: General, 130(1), 29–58. 10.1037/0096-3445.130.1.29

52. Zhou, D., Cornblath, E. J., Stiso, J., Teich, E. G., Dworkin, J. D., Blevins, A. S., & Bassett, D. S. (2020). Gender Diversity Statement and Code Notebook v1.0 (v1.0)[Computer software]. Zenodo. 10.5281/zenodo.3672110

53. Zuo, X., Honey, C. J., Barense, M. D., Crombie, D., Norman, K. A., Hasson, U., & Chen, J. (2020). Temporal integration of narrative information in a hippocampal amnesic patient. NeuroImage, 213, 116658. 10.1016/j.neuroimage.2020.116658

54. Zwaan, R. A. (1996). Processing narrative time shifts. *Journal of Experimental Psychology: Learning*, Memory, and Cognition, 22(5), 1196–1207. 10.1037/0278-7393.22.5.1196

55. Zwaan, R. A., Magliano, J. P., & Graesser, A. C. (1995). Dimensions of situation model construction in narrative comprehension. *Journal of Experimental Psychology: Learning*, Memory, and Cognition, 21(2), 386–397. 10.1037/0278-7393.21.2.386

